# Macro evolutionary patterns do not predict micro evolutionary trajectories

**DOI:** 10.64898/2026.07.10.737868

**Authors:** Chase C James, Suzana Gonçalves Leles, Hagen Buck-Wiese, Zachary C Landry, Elizabeth Morris, Dustin J. Marshall, Naomi M. Levine

## Abstract

As the world’s oceans change in response to climate change, phytoplankton communities will adapt to warmer, more stratified surface waters via plasticity, evolution, and range shifts. Current global ocean models assume that size structured phytoplankton communities have fixed trait relationships, and as a result generally predict that smaller size classes will become more dominant globally. However, this general expectation fails to consider how intra-species trait tradeoffs may operate orthogonally from large-scale inter-species tradeoffs—allowing for alternative evolutionary pathways given the limits and/or possibilities available to ancestral populations. To identify evolutionary pathways phytoplankton populations might take, we develop a novel modeling framework that combines a trait-based phytoplankton quota model with stochastic evolution (ecoTRACE). EcoTRACE explicitly decouples key phytoplankton traits from interspecific allometric relationships, allowing for novel phenotypes to emerge. We validated ecoTRACE against a long-term artificial size selection experiment on Dunaliella tertiolecta. We show that ecoTRACE captures multi-dimensional evolved phenotypes that quota models based on interspecific relationships fail to reproduce. Under fluctuating multi-stressor growth, model populations evolve phenotypic plasticity that deviates from predicted interspecific allometric relationships. EcoTRACE provides a framework for generating hypotheses as to the evolutionary trajectories that phytoplankton will experience in a warmer, more variable ocean.

## Introduction

As the climate warms, the ocean environment is rapidly changing with increased surface temperatures, stronger stratification, reduced nutrient supply to the surface ocean, higher oxidative stress, and potentially higher rates of variability across the globe (Oliver et al. 2018, Cheng et al. 2020, Cai et al. 2022). These changes are resulting in novel multi-stressor environments that will exert selective pressure on marine organisms to adapt to this new world (Collins et al. 2022). Marine microbes are the invisible workforce in the oceans driving biogeochemical cycles and supporting the ocean foodweb. With short generation times, the rate of change in the ocean environment matches the adaptation timescales for these organisms (Richter et al. 2022). Understanding and improving our predictions with regard to how marine microbes will adapt is therefore critical if we want to understand how carbon cycling and other global-scale ecological processes will be altered as a result of climate change.

Here we focus on photosynthetic marine microbes (phytoplankton), which form the base of pelagic food webs and mediate the transfer of carbon through the biological pump at regional and global scales. Current, state-of-the-art global-scale ocean models, such as those used to predict future ocean states, represent marine phytoplankton communities with a finite number of functional groups. A common approach is to define the traits associated with the different functional groups using empirically derived interspecific allometric relationships between key phytoplankton traits (such as nutrient uptake rates or maximum photosynthetic rate, P_max_, Figure 1a-b). These models are able to recover the current day biogeographical patterns and processes thought to shape these communities on a global scale (Follows et al. 2007, Stock et al. 2014, Aumont et al. 2015). Based on their success at capturing present day communities, these models have also been used to explore how phytoplankton distributions, diversity, and ecosystem structure may respond to future climate change (Bopp et al. 2013, Stock et al. 2014, Henson et al. 2021, Anderson et al. 2023). However, these future predictions implicitly inherit a key assumption that inter-species allometries, driven by millions of years of speciation and evolution, can be used to predict the adaptation trajectories present-day phytoplankton are likely to take in response to climate change: or more simply the assumption that macro-evolutionary patterns predict micro-evolutionary outcomes.

**Figure 1:**
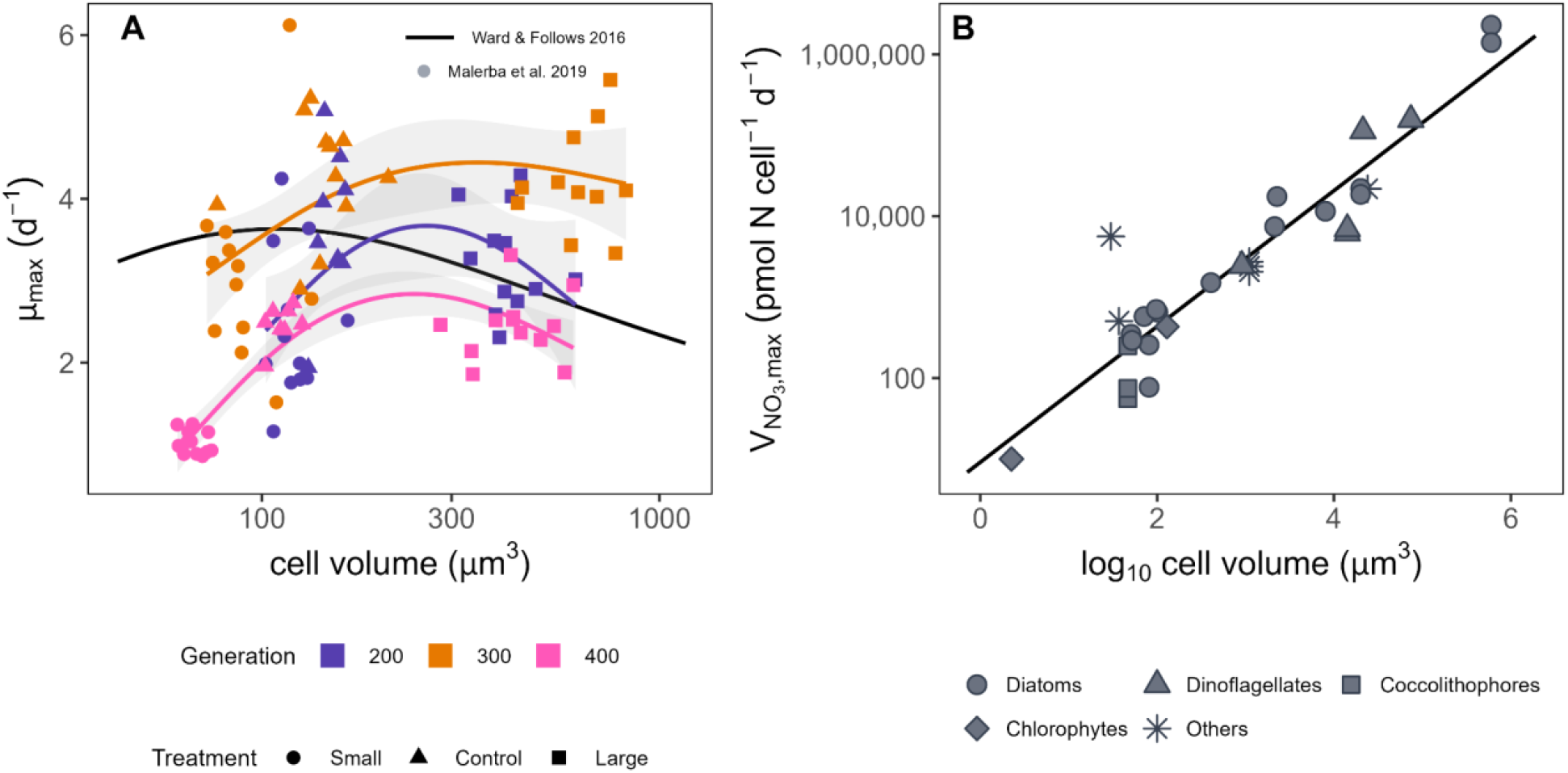
**A**. Data on variable growth rates across size-selected lineages of *Dunaliella tertiolecta* and the variation in size-growth rate allometric patterns across generations from Malerba and Marshall (2019) plotted against the solid line representing the unimodal allometry used in Ward and Follows (2016). **B**. Replotting of data from Edwards et al. (2015) highlighting the interspecies relationship between cell volume and V_NO3,max_ for marine phytoplankton species. Line represents a generalized linear model fit to this data.

While interspecific relationships can provide useful constraints on geological timescales, we argue that on shorter timescales intraspecific adaptation may not follow these relationships. There is increasing evidence that intraspecific trait variation and the mechanisms shaping these dynamics can produce responses that are distinct from, and sometimes directly opposite to, those observed across species (Agrawal 2020; Malerba et al. 2018; Brandenburg et al. 2018; Malerba and Marshall 2019; Levine et al. 2024). This likely arises, in part, because speciation introduces genetic tradeoffs whose associated costs and benefits differ from the broader processes characterized across species. In an artificial size-selection experiment on the green alga *D. tertiolecta*, Malerba et al. (2019) found that the relationship between cell size and growth rate varied across generations under strong selection, illustrating how adaptive responses within species can depart from expectations derived from interspecific patterns (Figure 1a).

Here we focus on size evolution because size is often considered a master trait in multi-class phytoplankton models such that most other traits are directly linked to cell size. Using these relationships to predict adaptation in models suggests that evolution is constrained along a single axis of trait variability. Moreover, while many biogeochemical models allow plasticity for some phytoplankton traits (primarily stoichiometry (N:C) and chlorophyll content (Chl:C)) these models typically do not allow for size plasticity in response to environmental conditions. This is inconsistent with decades of experimental evidence that phytoplankton cells exhibit large amounts of phenotypic plasticity especially in size (e.g. cells get smaller under nutrient stress (Peter and Sommer 2013), and temperature (O’Donnell et al. 2021)). Overall, phenotypic plasticity (both of cell size and other traits) is a fundamental mechanism that cells employ to mitigate environmental stress. This plasticity can also decouple cell phenotypes from the macro-evolutionary relationships employed by trait-based models.

To better describe how phytoplankton may adapt to a rapidly changing ocean, models must be able to correctly capture micro-evolutionary trajectories of multi-trait phenotypes. We show that by decoupling key processes from inter-specific allometric constraints we can better capture observed evolutionary trajectories. Critically, instead of relying on interspecific relationships, we explicitly incorporate cellular level trait trade-offs there by allowing additional phenotypic plasticity in cell size, stoichiometry, and chlorophyll content. In our new model framework, ecoTRACE, we embed a modified quota model within an eco-evolutionary framework allowing for multi-trait adaptation. We validate ecoTRACE against *D. tertiolecta* size-selection experiments (Malerba et al. 2018) to show that macro-evolutionary relationships do not predict the observed evolutionary trajectories. And finally, we show that departures from inter-specific allometries are not random but follow predictable evolutionary trajectories, wherein species evolve phenotypic plasticity in proportion to environmental variability.

## Results

### Representing Trait Correlation and Evolution in a quota model

#### New model framework

Allowing a quota model to capture phenotypic plasticity and trait adaptation requires a reformulation of the model. Here, we developed a model which combines **TRA**it **C**orrelation and **E**volution (Walworth et al. 2021) with ecological dynamics (**ecoTRACE**). The core biomass model in EcoTRACE is built on the backbone of previously developed quota models (Droop 1968, Geider et al. 1997, Ward et al. 2012) with several critical modifications which allows for additional decoupling between traits. Specifically, we link cellular metabolism to core space and energetic trade-offs rather than intraspecific trait relationships. These changes in model formulation allow traits to mutate freely in the model while providing mechanistic constraints on infeasible trait combinations (e.g. based on energetic constraints). Ultimately, this permits the emergence of novel phenotypes in the model as a function of the selection environment.

In EcoTRACE, phytoplankton can dynamically change their cell size. These cellular size changes are then mechanistically linked to other traits through shifts in cellular processes. For example, nutrient uptake is driven by the environmentally determined cell volume rather than a static size, respiration rates are split into specific mechanism-associated partitions, and maximum photosynthetic capacity is determined by total chlorophyll concentration rather than cell size (*see Methods*). We also leverage cellular trade-offs derived from a phytoplankton proteome model (Leles and Levine 2023) to constrain traits in the model. For example, we utilize an emergent allometry between the minimum Nitrogen:Carbon quota and size from the proteome allocation model (Supplemental Figure 1) which is based on intracellular space constraints and required cellular components (e.g. nucleus and ribosomes). We then validated this theoretical relationship with experimental data (Supplementary Figure 1).

Populations within EcoTRACE are represented as “super-individuals”, defined as a collection of cells that share all traits (i.e., identical parameters in the model, Figure 2a). The phenotype of the super-individual is represented using a quota model formulation (*described above and in Methods*) which estimates growth rates, mortality rates and the consumption of resources. Each model simulation represents hundreds of super-individuals which compete against each other for resources. As each super-individual grows in biomass, mutations occur and new super-individuals are generated (i.e. new parameter combinations which deviate slightly from the parent individual). Critically, in ecoTRACE, the model population modifies the resource environment and fitness is determined by competitive advantage (Figure 2b).

**Figure 2:**
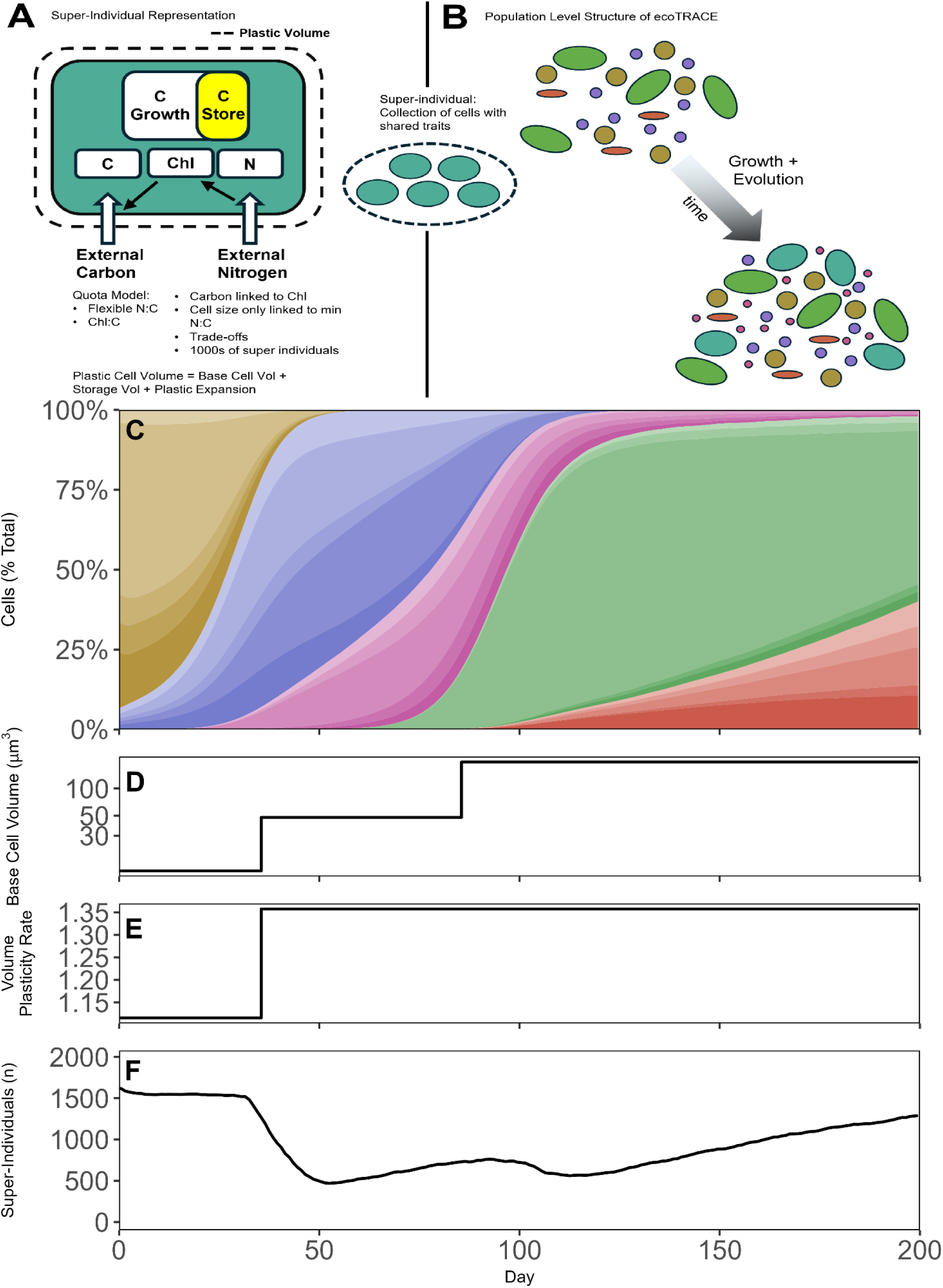
EcoTRACE: A high-resolution quota model nested within an eco-evolutionary framework. **A**. A cellular representation of the quota model, highlighting main pathways and parameters **B**. Schematic representation of ecoTRACE at the population level, showing multiple super-individuals within a population and how mutations can lead to a changing community composition over time **C-F:** Example ecoTRACE run showing how mutations emerge within a population **C**. Relative abundance of super-individuals, colors represent super-individuals binned by a combination of the two mutatable parameters (base cell volume and expansion rate) **D**. Base cell volume (μm^3^) of the most abundant super-individuals over time **E**. Expansion rate of the most abundant super-individuals over time **F**. Number of super-individuals over time.

General model dynamics are illustrated in Figure 2c-f. In this example simulation, we initialized a population in one environment (Environment A) before transitioning to a novel environment (Environment B). In Environment A, the gold genotype has the highest fitness but is quickly outcompeted in Environment B by genotypes with increasing levels of fitness in sequential selective sweeps (purple, then pink, then green, then red). With each selective sweep, we see a substantial shift in population level traits and a corresponding drop in diversity (Figure 2c-f). After several hundred generations, we observe stabilization both in the phenotype (traits) and in diversity as the population nears optimal trait values (Supplementary Figure 2). EcoTRACE is formulated such that it can be run to mimic lab experiments or natural environments and provides a testing ground to explore the interplay between intraspecies competition, plasticity and evolution under different and variable selection regimes.

#### EcoTRACE Validation

Validating a model of trait adaptation is challenging as it requires an extensive dataset with both trait evolution over time and the necessary metadata to correctly simulate the selective pressures imposed on the organism. Here, we leverage a long-term selection experiment on *Dunaliella tertiolecta*, a fast growing, cosmopolitan green alga (Malerba et al. 2018). This experiment provides an ideal validation dataset because the authors quantified trait evolution over 1500 days under experimentally controlled size selection (Malerba et al. 2018, Malerba and Marshall 2019, Malerba et al. 2020). By the end of the experiment, the cell size for large lineages was over 10x bigger than the small lineages, providing a novel dataset for testing how adaptation within a single species may occur when faced with strong selective pressure. This study also allows us to assess the difference between inter- and intraspecific relationships between cell size and other traits. We ran the ecoTRACE model to simulate the experimental design of Malerba et al (2018) (*see Methods*) and compared both the evolution of size over time and the resulting phenotypes for each lineage (nitrogen uptake, chlorophyll content, C:N, carbon/cell, chlorophyll/cell, carrying capacity, cell size plasticity, Figure 3 and Supplementary Figure 3).

**Figure 3:**
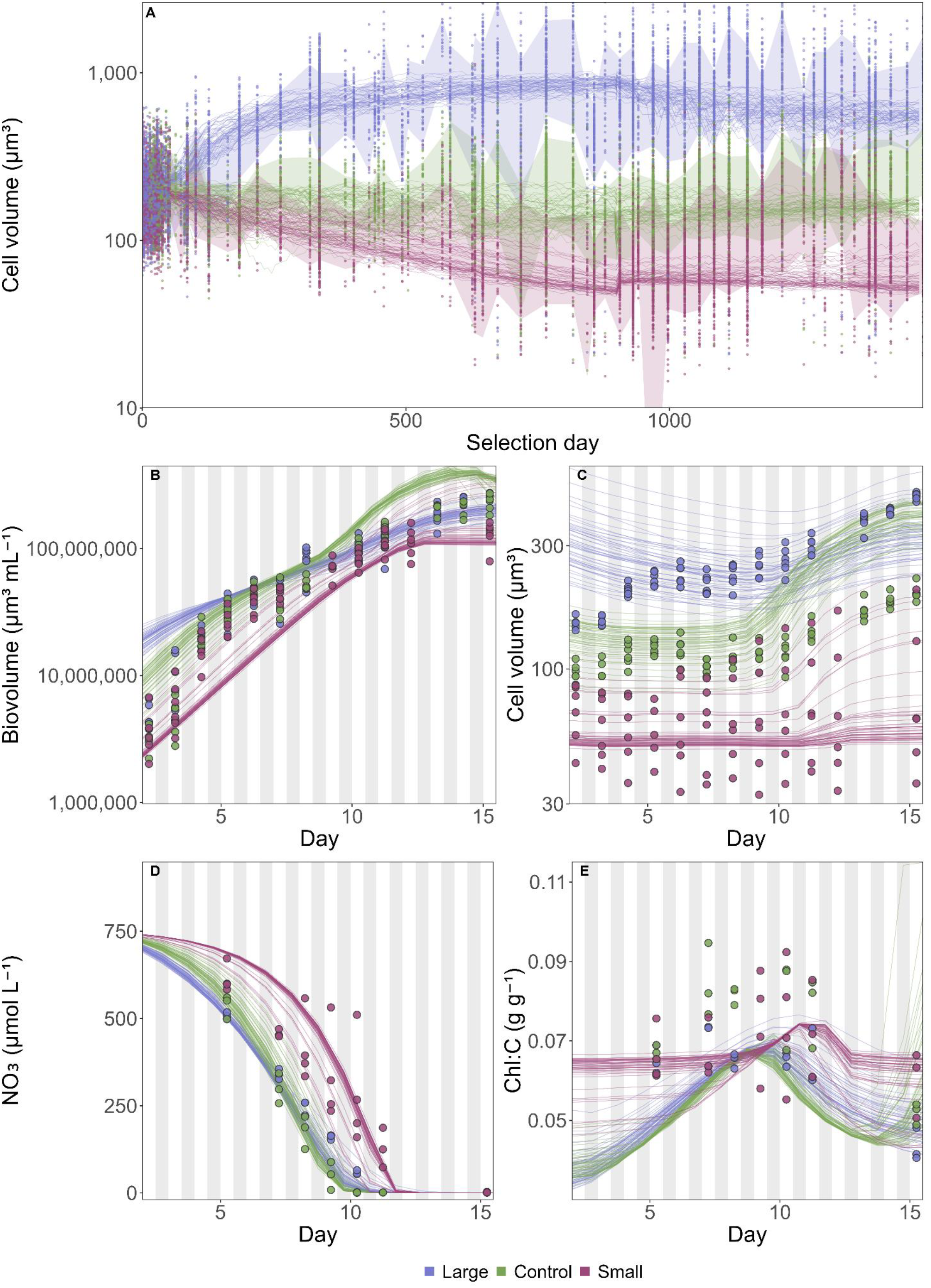
EcoTRACE validation against *D. tertiolecta* artificial size selection lineages. **A**. Validation comparing trajectories of cell volume (μm^3^) for artificial size selection experiments by the Marshall group and the ecoTRACE model. Points show a subset (10% random subset of the data) of actual cell volume measurements for the individual lineages within the experiment. Ribbons highlight ± 1 SD of cell volume over time per treatment (Large, Control, Small). Lines represent individual model runs (50 runs per treatment). The dashed vertical line denotes the switch in the size selection regime in the experiment from every 3-4 days to weekly (model runs reflect this change in the selection regime). **B**. Comparison between total biomass (um^3^ mL^-1^) in the model and growth curve experiments **C**. Comparison between cell volume (μm^3^) in the model and growth curve experiments **D**. Comparison between environmental NO_3_ (μmol L^-1^) in the model and growth curve experiments. **E**. Comparison between Chlorophyll:Carbon quotas (g g^-1^) in the model and growth curve experiments. Gray shading in **B-E** represents day night cycling.

EcoTRACE was able to capture the trajectories in cell size observed in the small, control, and large evolved lineages (Figure 2a). Later in the experiment, some small lineages appear to begin to slowly increase in size (despite strong selection pressure, *see Methods*). This is consistent with the hypothesis proposed by Malerba et al. (2018) that small lineages were nearing a boundary of size selection evolution for *D. tertiolecta*.

In addition to matching the observed changes in cell size over the course of the evolution experiment, the model also captured the realized multi-trait phenotype of the evolved lineages at the end of the experiment (Figure 2b-e). Across both the experimental data and the model, we identified convergence between lineages in total population biovolume (cell volume × cell count) over time (Figure 2b). The emergent model phenotype predicted substantial cell volume plasticity over the growth curve which matched the observations, with high cell volume plasticity in the large and control lineages and very little volume plasticity in the small lineages (Figure 2c). In the data and the model simulations, the small lineages showed the greatest degree of variability between lineages where the majority of lineages maintained a constant cell volume across the growth curve, but a few lineages increased cell size substantially during the stationary phase (Figure 2c). In both the model and the data, nitrogen uptake rate for small lineages was slower when compared to large and control lineages, which were more similar (Figure 2d). The model was also able to capture the observed changes in Chl:C across the growth curve with all lineages hitting their maximum Chl:C during mid to late exponential growth (Figure 2e). Finally, the model showed good agreement between observed total cells, total carbon, carbon per cell, and Nitrogen:Carbon over the growth curve for all lineages (Supplementary Figure 3). When taken together, the evolutionary trajectories coupled with the growth curve validation highlight the ability of ecoTRACE to not only capture the effect of artificial size selection but also capture the realized phenotypic plasticity both within and across evolutionary lineages.

#### Comparison between ecoTRACE and a quota model with inter-specific trait relationships

EcoTRACE is formulated to capture trait evolution with a framework that allows for multi-trait adaptation that can deviate from interspecific allometric relationships. However, the model does still link cell size to traits based on known energetic and space constraints on cellular metabolisms. Thus, it is important to ask two questions: 1) Can ecoTRACE capture well established relationships between environments and traits? And 2) How does the ecoTRACE formulation differ from more conventional quota model dynamics? We wish to stress that this analysis is not to suggest that inter-specific quota models are poor representations of reality. Rather, it highlights that these models were not designed to capture adaptive responses arising from the inherent multi-trait phenotypic plasticity of phytoplankton.

Theory predicts and experimental data confirms that cell size varies as a function of resource availability such that resource rich environments favor larger cells and resource depleted environments favor small cells. We tested the ability for ecoTRACE to capture this basic ecological principle through a set of simulations in which populations were allowed to evolve under constant high or low nutrient concentrations (akin to growth in a chemostat). EcoTRACE results align strongly with expected relationships where populations exposed to constant high nutrients evolved large, fixed cell sizes and populations exposed to constant low nutrients evolved small, fixed cell sizes (Figure 4a). We then repeated the simulations with increased levels of environmental variability. EcoTRACE predicted that cell size would vary plastically in response to the experienced environmental variability. Specifically, under episodic high nutrient conditions the model predicted substantial variation in both cell size and plasticity (Figure 4). Critically, under the 50- and 100-day nutrient pulses, which resulted in the highest degree of experienced environmental variability, populations diverged substantially from the inter-specific allometric relationship between cell volume and V_Nmax_ (Figure 4a) and between cell volume and nitrate uptake rates (V_NO3_, Figure 4b). In general, environmental variability with low nutrient conditions selected for small, minimally plastic cells, suggesting that total nutrient availability imposed a stronger selective pressure than the potential benefits associated with physiological plasticity.

**Figure 4:**
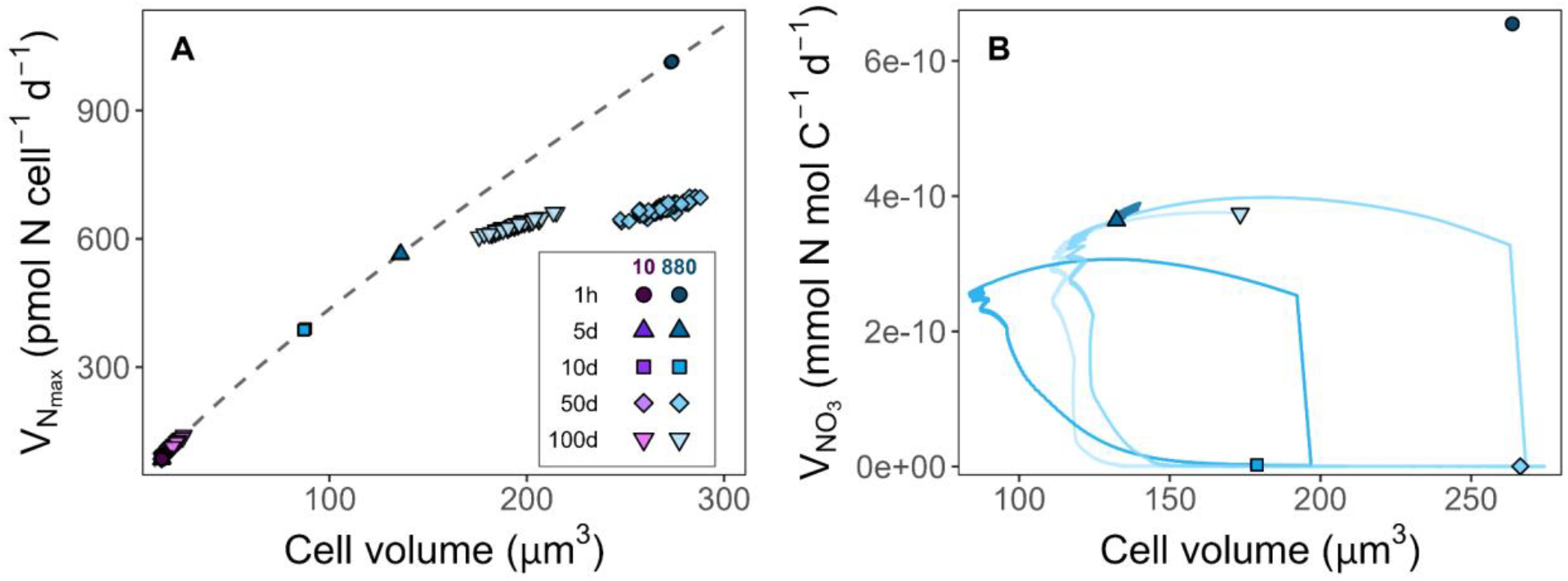
EcoTRACE dynamics under variable nutrient stress. **A**. Relationship between the median cell volume for the population and realized V_Nmax_ for the median cell volume given the mechanistic constraints to V_Nmax_ imposed on plastic cells. The dashed black line represents the inter-specific cell volume x V_Nmax_ allometry. **B**. Temporal changes in cell volume and nitrate update rates (V_NO3_) for representative runs over the last dilution period. Colors and shape identify each treatment type.

Combined, these simulations highlight how inter-specific allometries can still hold under equilibrium, to near-equilibrium conditions. However, populations quickly adapt, moving off inter-specific allometries as conditions become more variable.

Next we assessed how ecoTRACE predictions of trait adaptation differ from a conventional quota model formulation. We repeated the *in silico* representation of Malerba et al (2018) used to validate ecoTRACE (*above*) using a conventional quota model where interspecific allometric relationships were tied to the master-trait of cell size instead of the modified version of the quota model. Specifically, we combined ecoTRACE’s evolutionary framework with a modified version of the quota model from Ward and Follows (2016), see Methods for details.

Both ecoTRACE and the inter-specific quota model showed good agreement to the observed trajectories in cell-volume reported by Malerba et al. (2020), demonstrating that, with appropriate species-specific parameterization, inter-specific quota models can recover the evolutionary cell-size trajectories observed in the *D. tertiolecta* experiment (Figure 5a). However, examination of the resulting evolved phenotypes revealed a clear divergence between ecoTRACE and the inter-specific quota model predictions. Because the inter-specific model uses fixed allometric relationships and does not allow cell size plasticity, it was unable to accurately capture the evolved phenotypes within multidimensional trait space (Figure 5b-e and Supplementary Figure 4). Across all described traits (Total Biovolume, Cells, Total Carbon, cell volume, NO_3_ (media), Chl:C, and N:C) we observed better fits between model predictions and experimental observations from ecoTRACE relative to the inter-specific quota model (RMSE, Supplementary Table 1). Although both models achieved comparable accuracy in predicting total biovolume (RMSE; p > 0.05), the inter-specific quota model was unable to reproduce the magnitude and rate of biovolume change, yielding a predicted-versus-observed slope of 0.54 versus 0.94 for ecoTRACE. Both models showed weaker agreement with observed Nitrogen:Carbon ratios than for other response variables. However, this discrepancy likely reflects differences in the timing of nitrogen dynamics between model predictions and observations, particularly during the initial growth-curve spin-up phase, which were not explicitly represented in the validation framework to maintain model simplicity (*see Methods for details*).

**Figure 5:**
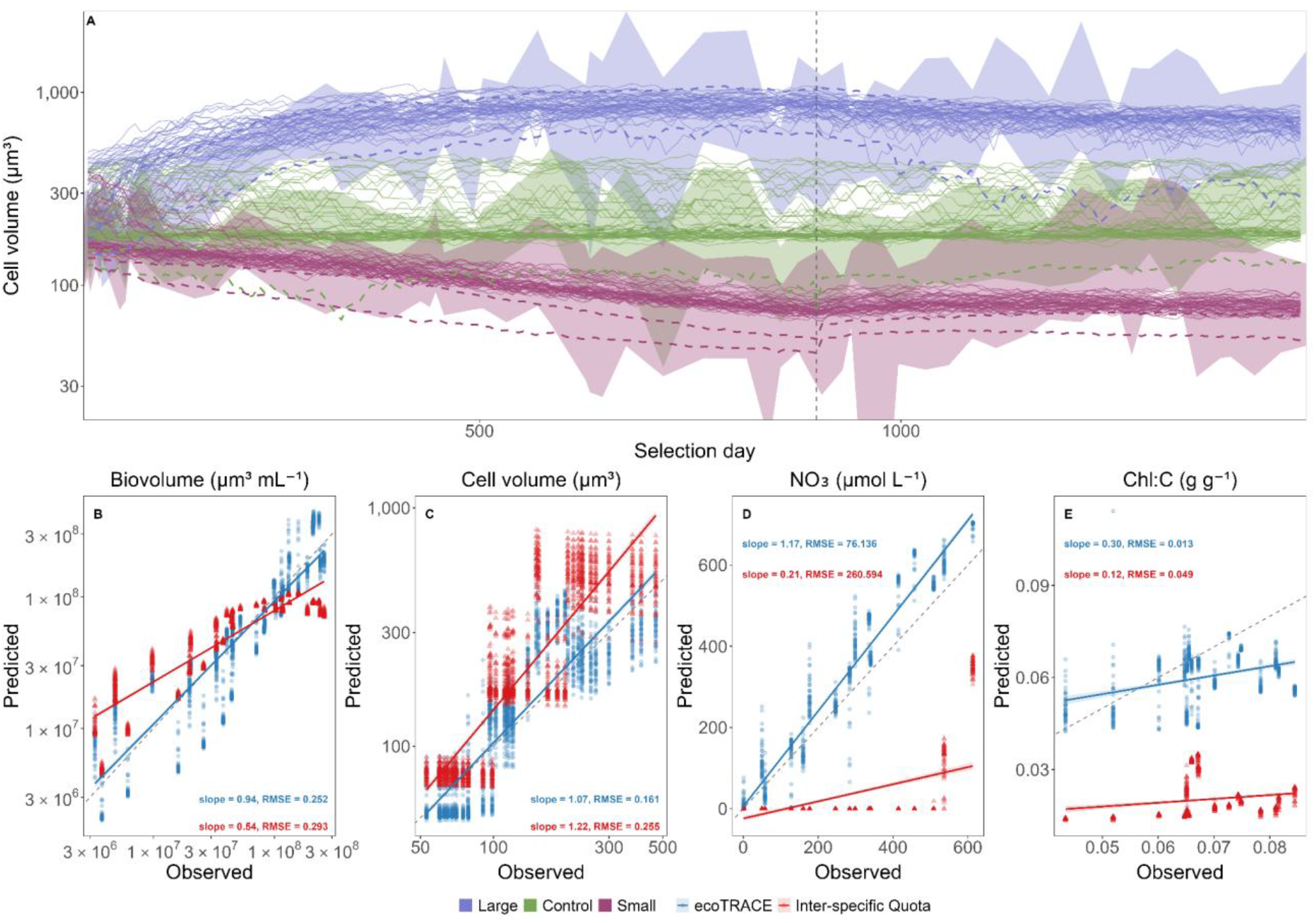
Comparison between observations and model predictions for the *D. tertiolecta* artificial size selection experiment. **A**. Validation comparing trajectories of cell volume (μm^3^) for artificial size selection experiments by the Marshall group, the ecoTRACE model, and the inter-specific quota model. Ribbons highlight ± 1 SD of cell volume over time per treatment (Large, Control, Small). Dashed colored lines represent the upper and lower margins of ecoTRACE model run trajectories per treatment (Large, Control, Small) shown in Figure 4a. Solid lines represent individual model runs for the inter-specific quota model (50 runs per treatment). The dashed vertical line denotes the switch in the size selection regime in the experiment from every 3-4 days to weekly (model runs reflect this change in the selection regime). **B-E** Observed (experimental) versus Predicted (either ecoTRACE or inter-specific quota) relationships in key phenotypes. Colored points indicate fit between model and experimental data. Lines represent a generalized linear model fit across all data per model. Slope and RMSE values are shown per model with colored text to indicate model **B**. Observed vs predicted total Biovolume (μm^3^ mL^-1^) **C**. Observed vs predicted cell volume (μm^3^) **D**. Observed vs predicted NO_3_ media concentration (μmol L^-1^) **E**. Observed vs predicted Chlorophyll:Carbon ratio (g g^-1^)

The current state-of-the-art quota models are powerful tools for providing context and accurate predictions about phytoplankton community composition and function in the present day. The conventional ecological models are also able to reproduce plausible evolutionary trajectories for a single trait. However, they struggle to accurately capture how species utilize their inherent physiological plasticity to adapt and evolve within a multidimensional trait space. Our results caution against extrapolating these inter-specific models to settings where adaptation selects on multi-trait plasticity thus leading to divergent evolved phenotypes on decadal to centennial timescales.

#### Exploring trait-space under multi-stressor selection

Above we focused on the impact of a single selective pressure (either nutrient availability or size selection) on the evolved multi-trait phenotype. In the environment, organisms are faced with complex environments such that organisms are often faced with conflicting trait trade-offs (Leles et al. 2025). Here we assess how phenotypic plasticity differs under multi-stressor environments (nutrients and light limitation) across a range of scales of variability for each stressor. We conducted an array of simulations with episodic high nitrate concentrations (5 days, 10 days, 50 days, and 100 day pulses) and variable light cycles (24 hr, 16 hr, 12 hr, and 6 hr light under both high light and low light intensities). Overall, we observe that low light selects for smaller cell sizes independent of nutrient regime (Figure 6a & 6b and light-only experiment Supplementary Figure 5) consistent with empirical evidence (Edwards 2015, Hillebrand et al. 2022). Variability in nutrient regimes drove larger phenotypic variations between cell volume and V_Nmax_ than differences in light availability, as expected (Figure 6a). Both nutrient and light variability resulted in multi-trait phenotypic plasticity in cell volume x N:C trait space (Figure 6b).

**Figure 6:**
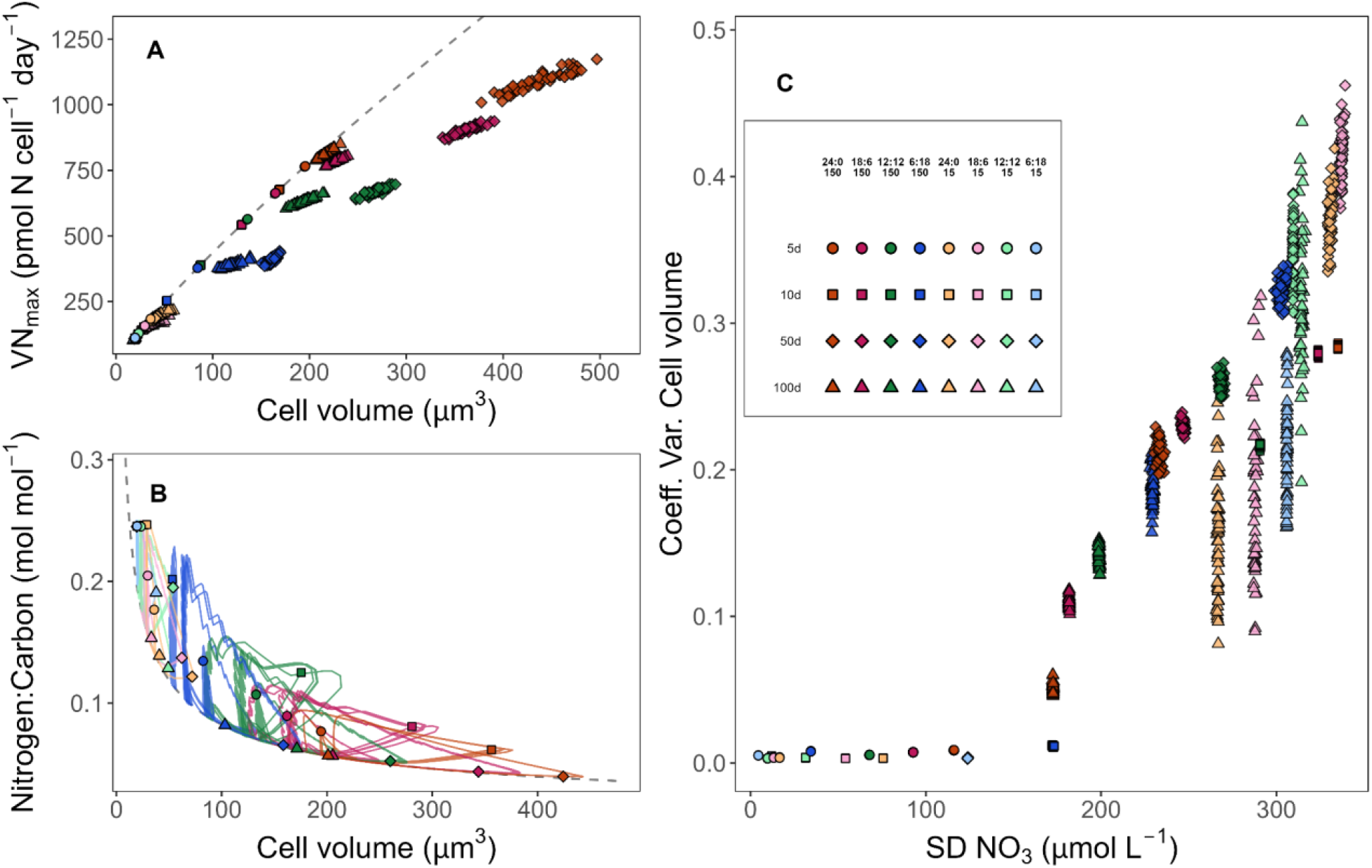
Effect of nutrient, light, and co-limitation on ecoTRACE evolutionary trajectories in both constant and variable environments. **A-C**. Results of combined nutrient-light co-limitation simulations across 32 environmental conditions. See panel **C** for corresponding color, tint, shape legend for each simulation **A**. Median cell volume versus V_Nmax_ across all simulations. **B**. Cell volume versus Nitrogen:Carbon ratio across all simulations **C**. Standard deviation of NO_3_ in the last 100 days of the simulation versus coefficient of variation (CV) of cell volume. Note within the legend the following: Colors (orange, red, green, blue) represent the main photoperiods within combination experiments. Tints represent the light level during the daytime period: 150 (orange, red, green, blue) vs 15 (yellow, pink, light green, light blue) W m^-2^. Shapes represent the timing between nutrient replenishments (5d, 10d, 50d, 100d).

Combined, these results further suggest that, under multi-stressor environments, cells do not converge on a single optimal allometric strategy but instead occupy a wide range of adaptive solutions shaped by interacting environmental constraints. EcoTRACE allows cell traits to exist plastically, however, this does not imply that cells optimize freely within this multidimensional trait space. We observed a strong positive relationship between variability in nutrient regimes (SD NO_3_) and variability in cell volume scaled to base cell volume (coefficient of variation of cell volume, Figure 6c) across all light intensities. Although lower light levels selected for smaller base cell volumes, these results suggest that populations evolve as much plasticity as is selectively advantageous for maximizing fitness under variable nutrient conditions, given the constraints imposed by the light environment.

## Discussion/Conclusion

A key challenge in marine biogeochemistry is to understand how marine microbial ecology will respond to a changing ocean and the implication for biogeochemical cycles. Addressing this requires integrating knowledge across the fields of biogeochemistry, microbial ecology, and evolutionary biology. Currently, trait-based quota models provide the best tool addressing these questions but modifications are needed to allow for phenotypic plasticity and trait evolution (Levine et al. 2024). The ecoTRACE model described above highlights the need to move away from interspecific allometric relationships and presents a framework through which multi-trait adaptation can be represented in biogeochemical models.

Macroevolution has resulted in clearly defined relationships between size and a variety of different traits for single celled organisms. These relationships provide a useful predictive framework for asking questions about present day communities. However, phenotypic plasticity is known to move traits off of interspecific relationships (Argyle et al. 2021, Malerba and Marshall 2019). Here we show that constraining model phenotypes to intraspecific allometries and forcing evolution to act along a single axis of trait variability is unable to capture observed evolved phenotypes. While it may be tempting to dismiss an artificial size-selection experiment as too removed from natural environmental selection to be broadly relevant, a core assumption embedded within trait-based models is that size functions as a master trait linking multiple core physiological tradeoffs. Consequently, in an experiment where selection acts explicitly on size, it is reasonable to expect that, if these interspecific tradeoffs are universally conserved within species, the model should accurately recover the resulting evolved phenotypes. The fact that it does not suggests that adaptive trajectories within a species may not be constrained by the same fixed allometric relationships that emerge across species. This is particularly important because traits such as cellular carbon content (Supplementary Figure 4c) may evolve through intraspecific adaptation in ways that are not predicted by models based on a single master variable, with potentially important consequences for forecasting future phytoplankton community structure and biogeochemical function.

As the ocean rapidly changes, we know the dial is “turning up” on many of the stressors experienced by phytoplankton communities. Temperatures are increasing, and enhanced stratification is likely to reduce nutrient availability in the surface ocean. At the same time, the magnitude and frequency of environmental variability is also projected to increase across many ocean regions (Oliver et al. 2018, Cheng et al. 2020, Cai et al. 2022). This shifting landscape requires a modeling framework capable of resolving adaptation within a multidimensional trait space. More broadly, these findings imply that accurately predicting future phytoplankton communities, and their associated biogeochemical functions, may depend on models capable of resolving adaptation as a process operating across multiple traits capable of decoupling from one another, rather than as a fixed extension of interspecific scaling relationships.

## Methods

### 5.1 EcoTRACE Model

EcoTRACE is a numerical model designed to simulate how a phytoplankton population grows, competes, and evolves over time. The model captures two distinct biological responses: 1) plastic phenotypic responses that are reversible changes in cellular traits driven by environmental changes (e.g. changes in cellular stoichiometry under nutrient stress); and 2) evolved responses which are heritable genetic changes driven by selection. As such, ecoTRACE is able to represent both short-term (acclimation) and long-term (adaptation) of cells to environmental change. The model tracks cell size as well as the internal state of each cell, specifically the carbon, nitrogen, and chlorophyll content (Figure 2a). In ecoTRACE, hundreds to thousands of genetically distinct cell types, called super-individuals, compete for resources within a shared environment. As cells grow and divide, random mutations can alter the heritable traits of daughter cells, allowing the population to evolve. Our model can thus be described as a trait-based quota model nested within a stochastic evolutionary framework, capable of tracking the population dynamics and internal regulation of large numbers of super-individuals, each with their own unique trait combinations. Below we first describe the quota model and then the ecoTRACE framework.

### Quota Model

Phytoplankton dynamics within ecoTRACE are represented using a trait-based quota model similar to Ward and Follows (2016) and Geider et al. (1997). Trait-based phytoplankton models define group specific traits using interspecific allometries. In ecoTRACE, these allometric relationships are used when there are strong mechanistic reasons to rely on these relationships (such as size versus maximum nitrogen uptake rate). However, for other traits, we decouple these relationships and specifically model metabolic mechanisms (such as total cellular photosynthesis) rather than rely on observed interspecies relationships. By not linking the cellular phenotype to a single master-trait, ecoTRACE allows for novel adaptations and phenotypes to arise. Three key features of ecoTRACE are: (i) cell volume plasticity responds to nutrient stress, (ii) carbon storage dynamics are explicitly partitioned between biosynthetic and storage pools, and (iii) total cellular photosynthesis is determined explicitly by the chlorophyll concentration within a given cell. EcoTRACE is formulated to explicitly resolve population dynamics over a diel-cycle. Here, we run with a 20-minute timestep.

### State Variables

As ecoTRACE is formulated to simulate the evolution of phytoplankton populations, each simulation is conducted with a large number of genotypes, hereafter referred to as super-individuals. Each super-individual, *j*, is associated with the following state variables: biomass carbon C_*bio,j*_ (mmol C cm^-3^), storage carbon C_*store,j*_ (mmol C cm^-3^), total carbon C_*j*_ = C_*bio,j*_ + C_*store,j*_ (mmol C cm^-3^), cellular nitrogen N_*j*_ (mmol N cm^-3^), cellular chlorophyll Chl_*j*_ (mg Chl cm^-3^), total cells Cells_*j*_ (Cells cm^-3^), base cell volume V_*base,j*_ (μm^3^), Expansion rate *ExpRate*_*j*_, and plastic volume V_*plastic,j*_ (μm^3^). Environmental state variables tracked within the model are the dissolved nitrate concentration (NO_3_, mmol cm^-3^), the dissolved ammonium concentration (NH_4_, mmol cm^- 3^), the incident irradiance (I_o_, W cm^-2^), and temperature (T, °C).

### Cell size

Each super-individual in the model is associated with a base cell volume V_*base,j*_. This is the minimum cellular volume allowed for that super-individual and sets key phenotypic traits.

Cellular carbon content (mmol cell^-1^) scales with base cell volume as (Supplementary Figure 6b):

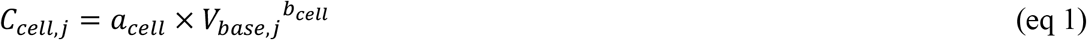

Where a_cell_ and b_cell_ are taxon-specific allometric coefficients (Table 1). Because this work focuses on simulations of *D. tertiolecta*, here we use *D. tertiolecta* specific values for a_cell_ and b_cell_.

**Table 1:** Parameters used in ecoTRACE model.

| Symbol | Parameter | Value | Units | Citation |
| --- | --- | --- | --- | --- |
| **Photosynthesis** |  |  |  |  |
| $P_{\text{Chl}}^{\text{max}}$ | Max Chl-specific photosynthesis rate | 72.86 | $\text{mmol C (mg Chl)}^{-1} \text{ d}^{-1}$ | Cullen 1990 |
| $I_{\text{sat}}$ | Light saturation irradiance | 0.0174 | $\text{W cm}^{-2}$ | Fitted (*D. tertiolecta*) |
| $a_s$ | Self-shading coefficient | 0.01036 | — | Fitted (*D. tertiolecta*) |
| $b_s$ | Self-shading exponent | -0.6127 | — | Fitted (*D. tertiolecta*) |
| **Quotas & Allometry** |  |  |  |  |
| $a_{\text{cell}}$ | C:V allometry coefficient | $3.71 \times 10^{-11}$ | — | Fitted (*D. tertiolecta*) |
| $b_{\text{cell}}$ | C:V allometry exponent | 1.130 | — | Fitted (*D. tertiolecta*) |
| $NC_{\text{max}}$ | Maximum N:C quota | 0.25 | $\text{mol N (mol C)}^{-1}$ | Fitted (*D. tertiolecta*) |
| $\text{ChlN}^{\text{max}}$ | Maximum Chl:N synthesis rate | 7.0 | $\text{mg Chl (mmol N)}^{-1}$ | Fitted (*D. tertiolecta*) |
| $NC_{\text{min}}$ | Minimum N:C quota (allometric) | $10^{-0.02523} \times V^{-0.52946}$ | $\text{mol N (mol C)}^{-1}$ | Leles & Levine (2023), proteome model |
| **NO<sub>3</sub> Uptake** |  |  |  |  |
| $a_{V, \text{NO}_3}$ | $V_{\text{max}}$ NO<sub>3</sub> allometry coefficient | $6.60 \times 10^{-12}$ | — | Edwards et al. 2015, adj. *D. tertiolecta* |
| $b_{V, \text{NO}_3}$ | $V_{\text{max}}$ NO<sub>3</sub> size exponent | 0.84 | — | Edwards et al. 2015, adj. *D. tertiolecta* |
| $a_{K, \text{NO}_3}$ | $k_{\text{NO}_3}$ allometry coefficient | $4.726 \times 10^{-6}$ | — | Edwards et al. 2015, adj. *D. tertiolecta* |
| $b_{K, \text{NO}_3}$ | $k_{\text{NO}_3}$ size exponent | 0.50 | — | Edwards et al. 2015, adj. *D. tertiolecta* |
| $\lambda_{\text{NH}_4}$ | NH<sub>4</sub> inhibition of NO<sub>3</sub> uptake | $4.6 \times 10^6$ | — | Ward & Follows 2016 |
| **NH<sub>4</sub> Uptake** |  |  |  |  |
| $a_{V, \text{NH}_4}$ | $V_{\text{max}}$ NH<sub>4</sub> allometry coefficient | $1.28 \times 10^{-11}$ | — | Edwards et al. 2015, adj. *D. tertiolecta* |
| $b_{V, \text{NH}_4}$ | $V_{\text{max}}$ NH<sub>4</sub> size exponent | 0.75 | — | Edwards et al. 2015, adj. *D. tertiolecta* |
| $a_{K, \text{NH}_4}$ | $k_{\text{NH}_4}$ allometry coefficient | $8.4 \times 10^{-8}$ | — | Edwards et al. 2015, adj. *D. tertiolecta* |
| $b_{K, \text{NH}_4}$ | $k_{\text{NH}_4}$ size exponent | 0.60 | — | Edwards et al. 2015, adj. *D. tertiolecta* |
| **Growth & Respiration** |  |  |  |  |
| $\beta_c$ | Biosynthesis cost | 0.466 | $\text{mmol C (mmol N)}^{-1}$ | Adjusted from W&F 2016 |
| $\text{basres}_{\text{ref}}$ | Basal respiration reference rate | 0.06 | $\text{d}^{-1}$ | Fitted (*D. tertiolecta*) |
| $V_{\text{ref}}$ | Basal respiration reference volume | 100 | $\mu\text{m}^3$ | Fitted (*D. tertiolecta*) |
| $\alpha_{\text{resp}}$ | Respiration allometric exponent | 0.80 | — | Fitted (*D. tertiolecta*) |
| $m$ | Linear mortality rate | 0.01 | $\text{d}^{-1}$ | Fitted (*D. tertiolecta*) |
| **Volume Plasticity** |  |  |  |  |
| $k_{\text{base}}$ | Minimum plasticity decay rate | 0.01 | $\text{d}^{-1}$ | Fitted (*D. tertiolecta*) |
| $k_{\text{max}}$ | Maximum plasticity decay rate | 3.25 | $\text{d}^{-1}$ | Fitted (*D. tertiolecta*) |
| $k_{s, \text{mult}}$ | Expansion $K_s$ multiplier | 1.5 | — | Fitted (*D. tertiolecta*) |
| $c_{\text{std}}$ | Standard plasticity respiration cost | 0.03 | — | Fitted (*D. tertiolecta*) |
| $c_{\text{exp}}$ | Expansion plasticity respiration cost | 0.02 | — | Fitted (*D. tertiolecta*) |
| **Evolution** |  |  |  |  |
| $p$ | Mutation probability per individual per division | 0.01 | — | Fitted (*D. tertiolecta*) |
| $\sigma_V$ | Mutation SD for base cell volume | 0.3 | — | Fitted (*D. tertiolecta*) |
| $\sigma_{\text{ExpRate}}$ | Mutation SD for expansion rate | 0.4 | — | Fitted (*D. tertiolecta*) |

Base cell volume is also used to set minimum values for basal respiration (R_basres_, eq 23-25), nitrogen uptake (VN_max_, eq 9-12,14,15,17), and nitrogen affinity (k_N_, eq 13 & 16). Plasticity in cell volume can then modify respiration and nitrogen uptake traits as described below.

In ecoTRACE, the base cell volume evolves to best match the average growth environment experienced by the cells such that cells which regularly experience high nutrient concentrations evolve large base cell sizes while cells in chronically nutrient limited environments evolve small cell volumes (*see Results*).

In response to environmental variability, super-individuals are also able to plastically expand their cell volume above V_*base,j*_ to temporarily increase their total surface area. This increases VN_max,j_ when external nutrients begin to decrease and allows cells to temporarily take up additional nitrogen, but only if internal nitrogen reserves are sufficient to support the metabolic cost of expansion. Cell volume plastically changes (μm^3^ day^-1^) as:

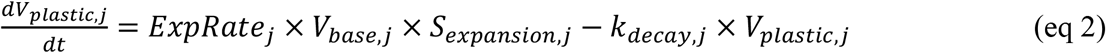

Where the expansion signal S_expansion_ is calculated as:

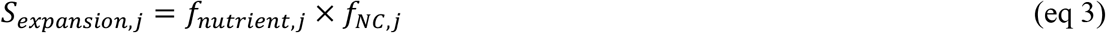

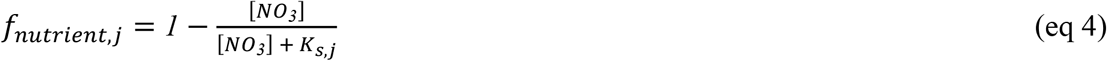

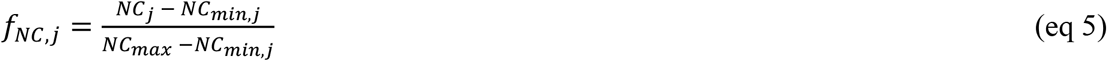

Where *NC*_*j*_ is the nitrogen-to-carbon content of the super-individual, *NC*_*min,j*_ is the minimum NC content, and *NC*_*max,j*_ is the maximum NC content. When nutrients are replete, f_nutreint,j_ = 0 and cells shrink back to their base cell volume (ΔV_plastic,j_ ≤ 0). Similarly, when the cellular quota is equal to NC_min,j_, f_NC,j_ = 0 and cells are not able to expand. The decay rate k_decay,j_ allows cells to shrink back to their base volume. This shrinkage rate is environment-dependent such that the k_decay,j_ is low under sustained nutrient stress and is elevated when nutrients are replenished:

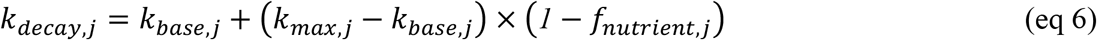

This formulation ensures that plastic volume is maintained under persistent nutrient limitation but rapidly relaxes when nutrients are restored. This produces the phenotype of larger cells in stationary phase which is consistent with both *D. tertiolecta* physiology (this study) and many other phytoplankton species (Lomas et al. 2024, Supplementary Figure 7).

The instantaneous cellular volume (μm^3^) for each super-individual (V_j_) is then calculated as:

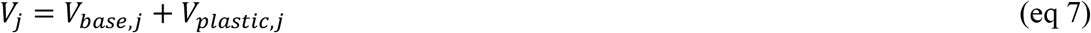

### Temperature Dependence

We assume temperature dependence on all cellular processes (e.g. nutrient uptake and carbon fixation rates) modeled using Arrhenius scaling (Li 1980, Geider et al. 1997)

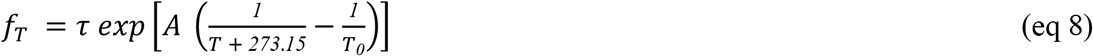

Where τ = 1.0 is a scaling constant, *A* = -4000K is the activation parameter, T_0_ = 298.15 K (25 °C) is the reference temperature, and T is in °C.

### Nitrogen uptake

Nitrogen uptake by cells follows Michaelis-Menten kinetics where nitrate 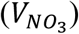 and ammonium uptake 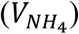 by cells in mmol N cell^-1^ d^-1^ are calculated as:

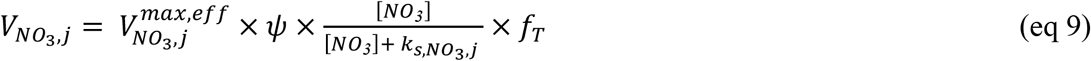

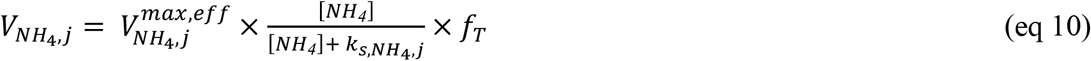

Where ammonium inhibition of nitrate uptake (ψ) is calculated as:

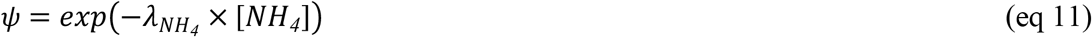

Where λ_NH4_ is a constant (Table 1).

Consistent with theory and observations, both maximum nitrate uptake rate 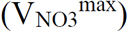 and half-saturation constant (k_s,NO3_) scale with cell size (Edwards et al. 2015). But here we modify the conventional size-based allometry to account for cellular plasticity. Specifically, we assume that the base cell volume sets 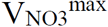 (mmol NO_3_ cell^-1^ day^-1^) and k_s,NO3_ (mmol cm^-3^):

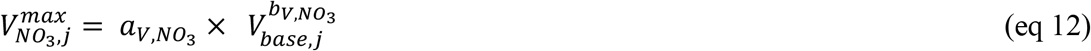

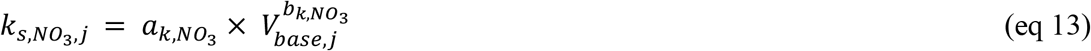

The values for a_V,NO3_ and b_V,NO3_ and a_k,NO3_ and b_k,NO3_ were derived from empirical interspecies allometries (Edwards et al. 2015). Here we modify this relationship by specifically adjusting the intercept for V_NO3_^max^ to match the subset of *D. tertiolecta* data present in Edwards et al. 2015. Both k_s_ for NO_3_ and NH_4_ were adjusted to reflect uptake kinetics observed in our growth curve experiments (Figure 3d, Table 1).

We assume that as a cell plastically expands in size that both V_Nmax_ and k_s_ increase. However, because the cell is plastically expanding we assume that the effective maximum rate (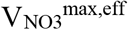, mmol NO_3_ cell^-1^ day^-1^) saturates via a ρ-dampened power law:

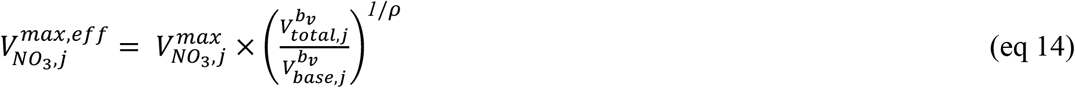

Where,

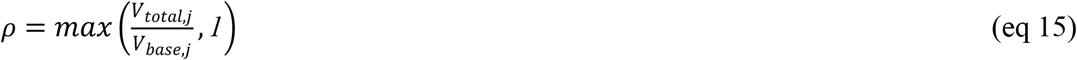

Similarly, the half-saturation constant (k_s eff,NO3_, mmol cm^-3^) grows at a super-linear rate (lower affinity per unit volume) as:

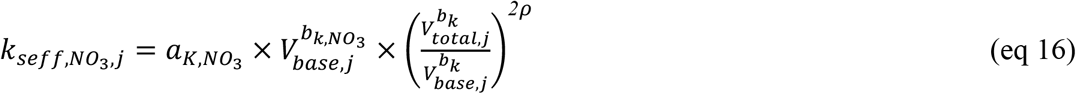

The same structure applies to 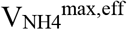 and k_seff,NH4_. This allows the model to differentiate between a cell that has plastically expanded to a given size from one that has a base cell volume of that size.

Finally, total nutrient uptake (VN_total,j_ = V_NO3,j_ + V_NH4,j_, mmol NO_3_cell^-1^ day^-1^) is constrained by:

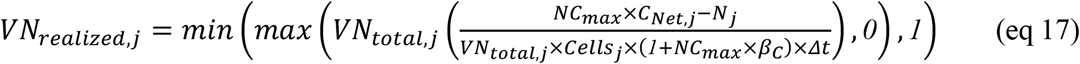

Where C_Net,j_ = P_C,j_ -R_basres,j_ x C_j_ (eq 26 excluding mortality, described below), and β_C_ is the biosynthesis cost (described below, Table 1). Eq 17 limits cellular nitrogen uptake such that cells do not exceed the maximum allowed nitrogen-to-carbon content of a cell (*NC*_*max*_).

We assume an allometric relationship between the minimum nitrogen-to-carbon content of cells (*NC*_*min,j*_, mmol N mmol C^-1^) and cell volume (μm^3^). Here we derive the relationship between *NC*_*min,j*_ and cell size using the proteome allocation model from Leles et al. (2023) and confirmed with experimental data (Supplementary Figure 1). Specifically, *NC*_*min,j*_ is calculated as:

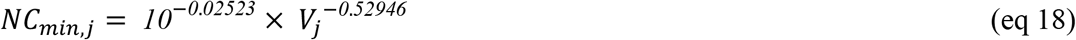

All cells have a fixed maximum: *NC*_*max*_ = 0.25.

### Photosynthesis

Carbon fixation by each super-individual (P_C,j_, d^-1^) is calculated as:

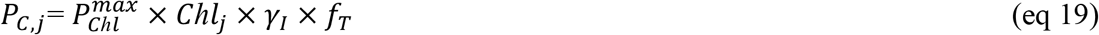

Where Chl_j_ is the cellular chlorophyll content of super-individual j, and 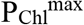 is the maximum per-chlorophyll photosynthetic rate in mmol C mg Chl^-1^ day^-1^ (Table 1). Light limitation (γ_I_) is calculated as:

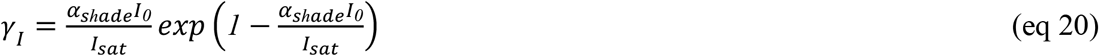

Where *I*_*0*_ is the incident light irradiance in W cm^-2^, *I*_*sat*_ is the irradiance at which 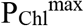 is released from light limitation, and α_shade_ accounts for the reduction in light due to community self-shading. α_shade_ is calculated as:

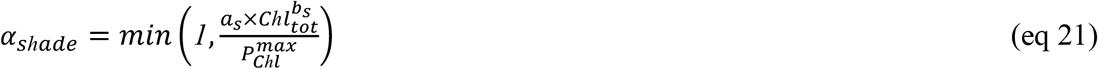

Where,

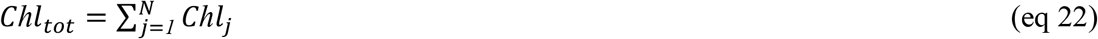

such that shading is dependent on the total concentration across all super-individuals. a_s_ and b_s_ are constants derived from experimental and environmental data (Table 1, Supplementary Figure 6a).

### Respiration

EcoTRACE partitions respiration into two categories: basal respiration and respiration due to metabolic biosynthesis. Basal respiration (d^-1^) is calculated as:

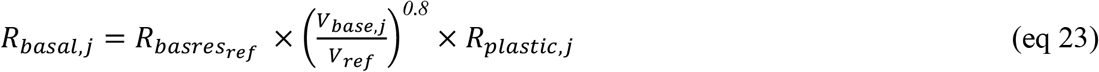

Where *basres*_*ref*_ and V_ref_ are constants (Table 1). R_plastic,j_ (d^-1^) represents the metabolic cost of maintaining plastic machinery and is calculated as:

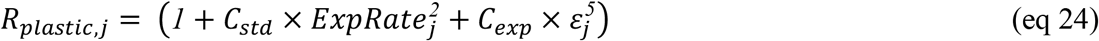

where C_std_ and C_exp_ are constants, and ε_j_ = (V_plastic,j_ / V_base,j_). C_std_ x ExpRate_j_^2^ imposes a penalty for mutations not currently undergoing conditions relevant to adaptive plasticity, stopping runaway mutations occurring during stable nutrient conditions. 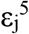 imposes a sharp, nonlinear penalty only when V_plastic,j_ is a large fraction of V_base,j_, penalizing cells that are currently expanded far beyond their base volume.

Respiration as the result of metabolic biosynthesis (d^-1^) is calculated based on the framework of Ward & Follows (2016) as:

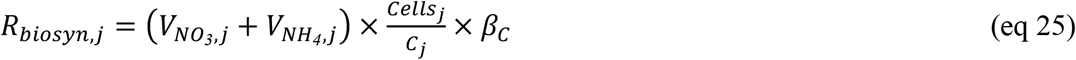

Where β_C_ (mmol C mmol N^-1^) is the carbon cost of nitrogen assimilation (Table 1).

### Mortality

A linear mortality rate (m = 0.01, d^-1^) is applied as a proportional loss to all state variables at each timestep.

### Biomass growth

The change in biomass 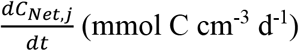of each super-individual is calculated as:

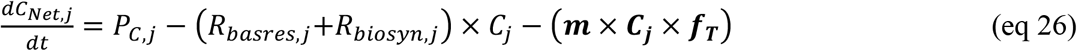

Where the first term (*P*_*C,j*_) is new carbon from photosynthesis, the second term is the loss of carbon due to respiration, and the third term is the loss of carbon due to mortality. To ensure that the nitrogen-to-carbon content of the super-individuals does not fall below their NC_min,j_ (eq 18), 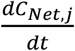is capped based on the realized uptake of nitrogen (*N*_*j*_ + *VN*_*realized,j*_):

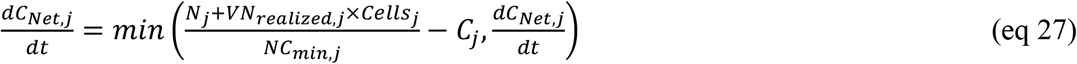

Thus, when cells hit *NC*_*min,j*_, any excess fixed carbon is released (excreted) rather than incorporated into biomass.

### Carbon Storage

Cellular carbon is divided into a biosynthetic pool (C_bio,j_, mmol C cm^-3^ d^-1^) and a storage pool (C_store,j_, mmol C cm^-3^ d^-1^). During the light period, photosynthetically fixed carbon is allocated first to biosynthesis and then to storage once the biosynthetic demand is met. During the dark period, stored carbon supports basal respiration, preventing the biosynthetic pool from being depleted. This explicit partitioning is critical for the sub-daily model resolution because it allows cells to carry excess carbon through the night without underestimating daytime photosynthetic capacity.

During the light period (C_Net,j_ > 0), carbon is split between storage and biosynthesis in proportion to the anticipated nighttime respiration demand:

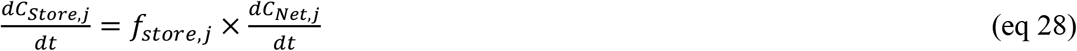

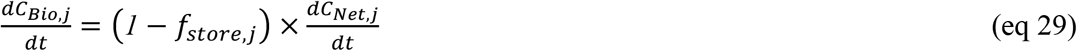

Where f_store,j_ is computed from the previous night’s total respiration load scaled to the current carbon pool. This approach was based on findings by Müller et al. (2026), which showed that diatoms store carbon in anticipation of nighttime demands, based on the demands of the previous night.

During the dark period and or when P_C,j_ < R_total,j_ (C_Net,j_ < 0), the respiration deficit is drawn from storage first (eq 30). Only if C_store,j_ is fully depleted is the remainder taken from C_bio,j_ (eq 31).

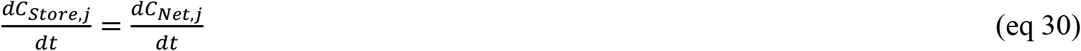

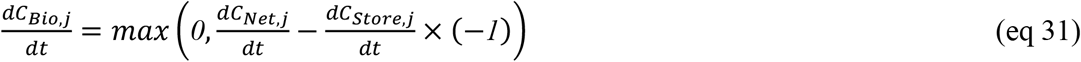

Change in cell count (cells cm^-3^ d^-1^) is derived at each step from biomass carbon:

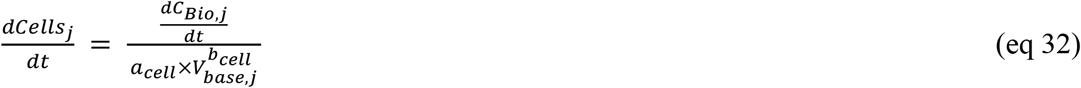

### Change in cellular nitrogen

The change in cellular nitrogen content (mmol N cm^-3^ d^-1^) of each super-individual 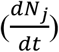is calculated as:

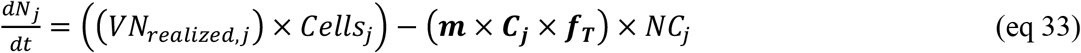

### Change in cellular chlorophyll

The change in chlorophyll content per cell 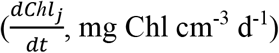is calculated as:

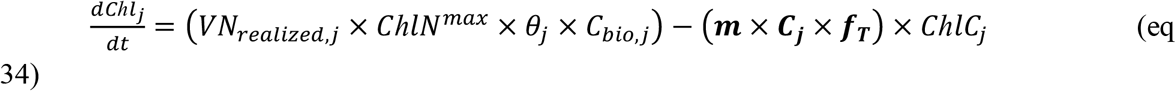

where *ChlN*^*max*^ (mg Chl mmol N^-1^) is the maximum Chlorophyll:Nitrogen synthesis rate (Table 1) and θ_j_ is the chlorophyll allocation factor (θ_j_), which balances light demand and nitrogen status:

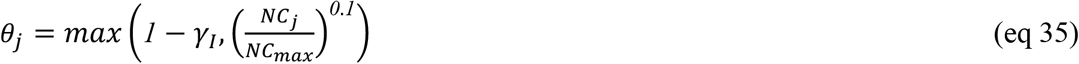

*θ*_*j*_ maximizes chlorophyll synthesis when cells are either light-limited, (γ_I_ < 1) thereby enhancing photon-capture, or are nitrogen-replete (*NC*_*j*_ near NC_max_).

### 5.2 Quota Model with inter-specific trait relationships

To contrast the new ecoTRACE quota model formulation against a model that used more inter-specific allo metric relationships to define phytoplankton traits, we implemented an alternative version of the quota model in ecoTRACE based on the core formulations of Ward and Follows (2016), hereafter referred to as the “inter-specific quota model”.

The inter-specific quota model defines maximum nitrogen uptake rate (VN_max_), half-saturation constant (K_s_), and maximum photosynthetic capacity (P_max_) for each super-individual based on inter-specific allometric relationships. This intrinsically links cell volume to many phenotypic traits such that, as cell volume evolves, all linked traits shift deterministically along prescribed inter-specific relationships.

Here, we used the formulations from Ward and Follows (2016) which do allow for plasticity in cellular C:N quota and chlorophyll content. We have simplified the Ward and Follows model to only include photosynthesis (no mixotrophy) and made nitrogen (NO_3_ and NH_4_) the only limiting nutrient within the model. We slightly modify the light limitation formulation to be consistent with the ecoTRACE quota model such that Ward and Follows S14 is the same but *I* is replaced by (α_shade_ x *I*_*0*_ / *I*_*sat*_). The remaining equations within the inter-specific quota model are directly comparable to the formulations present in Ward & Follows (2016).

The inter-specific quota model was run with a daily timestep (compared to ecoTRACE’s 20-minute timestep), consistent with the original Ward and Follows (2016) implementation. Because the model operates at daily resolution, photosynthesis and respiration are computed as daily averages and explicit carbon storage dynamics and diel cycling are not represented.

### Modifications for D. tertiolecta simulations

The inter-specific Ward and Follows (2016) parameterization is designed for the marine phytoplankton and was not derived for *D. tertiolecta* specifically. To ensure a fair comparison with the experimental data and with ecoTRACE, we applied the *D. tertiolecta* specific tuning for carbon-volume allometry and optimum temperature.

Key parameters that were not modified from the Ward and Follows (2016) values are listed in Supplementary Table 3.

### 5.3 Evolutionary framework

Each super-individual is defined by a genotype (parameters) with an emergent phenotype (e.g., *C*_*j*_, *N*_*j*_, *Chl*_*j*_, *V*_*plastic,j*_) which is governed by the quota model dynamics described above. Here, we allow two parameters to evolve, though ecoTRACE is designed to allow any parameter to mutate; 1) base cell volume *V*_*base,j*_ (μm^3^) and 2) expansion rate *ExpRate*_*j*_. For simulations with the inter-specific quota model only cell volume is mutated.

Mutations arise as the super-individuals grow. Specifically, when a super-individual accumulates enough cells to produce a daughter super-individual (defined here as five cells), it becomes eligible for mutation. At each timestep, all super-individuals that meet the ‘new daughter cell count’ threshold are candidates for new mutations. If a super-individual grows enough to generate multiple new daughter super-individuals, each new daughter is assessed separately for possible mutations. We determine whether the new daughter super-individual has experienced a mutation by drawing from a binomial distribution with a mutation probability (p, Table 1). If the daughter super-individual is determined to not have mutated, the biomass is added back to the parent cell and can not be mutated again until additional growth has occurred.

If a mutation has occurred, a new super-individual is created with a new set of parameters mutated from the parents parameters. We allow for new super-individuals to have mutations in either one or multiple parameters (here base cell volume only, expansion rate only, or both base cell volume and expansion rate). New traits are selected from a gaussian distribution where μ is the given trait value of the parent and σ is the mutation variance for that trait. This results in most new super-individuals having traits that are only slightly different from the parent values with infrequent cases with large changes (mutations) in trait value. We allow traits to both increase and decrease with mutation but impose a lower bound on the expansion rate at zero and minimum cell volume at 1 μm^3^. Mutation parameters are given in Table 1. Since the allometric relationships linking cell size to traits are log-linear, mutations to base cell volume are calculated in log_10_ space.

At creation, each new super-individual is given the identical phenotype as its parent (e.g. *C*_*j*_, *N*_*j*_, *Chl*_*j*_) and the associated biomass is subtracted from the parent’s pools. To avoid an exponentially increasing number of super-individuals, when the number of cells within a super-individual falls below a critical threshold (here set to one cell) it is assumed to be extinct and is no longer tracked in the model.

### 5.4 Model simulations

Below we describe the different model simulations presented in this paper. Due to the stochastic nature of ecoTRACE, each simulation type was conducted 50 times to capture both the mean response and variability in model dynamics.

### Size-selection simulations

Prior to running the size-selection experiments, the model was spun-up for 2,000 days under the experimental conditions used by Malerba et al. 2018. The models were initialized with a single super-individual with a base cell volume of 150 m^3^ and an ExpRate of 0.1. The model populations were diluted to 30,000 cells mL^-1^ every 7 days and NO_3_ was simultaneously replenished to 880 μmol L^-1^. Light (150 W m^-2^) was provided on a 14:10hr light:dark cycle and temperature was held constant at 25 °C. To account for stochasticity in the model simulations, 50 replicates of the spin-up were conducted. After 2,000 days, the models reached a steady state solution with an average population of ~2,000 super-individuals and an average cell size of 261 μm^3^ (range 167–355 μm^3^ over 50 replicates), which was consistent with the observed ancestral cell volumes which averaged 224 μm^3^ (with a range of 140-308 μm^3^). Following the spinup, each of the 50 replicates were run under all three treatment conditions (small, control, and large), such that each replicate represents a possible experimental run, from a shared ancestor through the selection simulation.

Simulations were then conducted to mimic the size-selection experiment of Malerba et al. (2018). In the experiment, size selection was done via centrifugation with the objective of a 10% change in mean cell volume across the population per selection event. Large cells were preferentially retained in the pellet and small cells in the effluent. The probability of survival for any individual in both the experiment and model is a function of the population characteristics at the time of selection. Specifically, both the distribution of cell sizes and the abundance of each cell type will impact the selective pressure on the individuals within the population. We represent this in the model using both an assessment of the relative size of the super-individual and the abundance of that individual in the population.

In the model, we represented this size selection through a percent shift in the mean cell volume across pre- and post-selection populations. During each size-selection event, super-individuals are given a transfer or survival probability based on their cell size relative to the mean cell size across all super-individuals in the population. Specifically, the relative cell size for each super-individual is computed as the biomass normalized z-score (z_j_) in log_10_ cell volume space as:

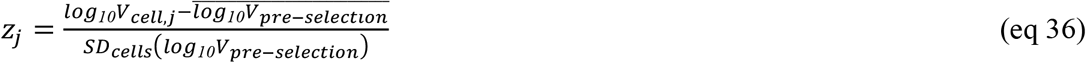

Where,

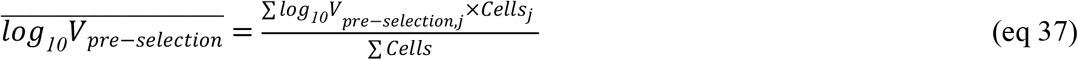

And,

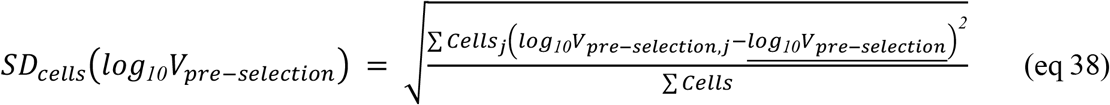

and the survival probability (p_j_) is calculated as:

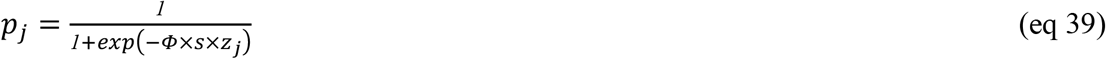

where s is selection (+1 for large, -1 for small) and Φ represents the steepness of the survivability curve. Specifically, Φ determines how likely a super-individual is to survive size selection given its relative size compared to the distribution of cell sizes within the population. When Φ = 0, cells are equally as likely to survive or die via size selection. Large values of Φ result in increased survivability for individuals that are larger than the mean (under large-size selection) or smaller than the mean (under small-size selection).

Take an example of two super-individuals under large size selection. Super-individual A is larger than the mean with a p_j_ of 0.75 and super-individual B is smaller than the mean with a p_j_ of 0.25. During size selection, 75% of super-individual A’s cells would be retained and 25% of super-individual B’s cells would be retained. Thus, if both super-individuals have equal population sizes pre-selection, then following size selection super-individual A would be three times more abundant than super-individual B and the total population size would be 50% less than before selection. Following size-selection, model populations are then further diluted to a constant biomass (3.125 μmol C cm^-3^). Dilution based on biomass rather than cell count was done for consistency with the experimental methodology of Malerba et al. (2018).

Experimentally, Malerba et al. 2018 adjusted centrifugation speeds and times continually throughout the experiment to achieve a constant selective pressure of ~10%. This was necessary due to the continually evolving populations which require different experimental procedures to accomplish the same selective pressure. Similarly, in the model we adjust the steepness of the survivability curve (Φ) throughout the simulation to account for shifts in the distribution and relative abundances of cell sizes in the model population. Specifically, at each size-selection, the value of Φ that provides the desired selective pressure (Ω) is determined where:

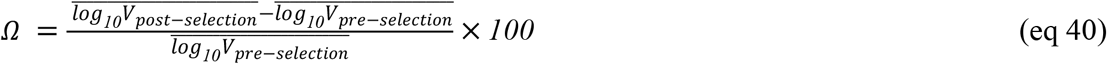

This is akin to changing the centrifugation protocol as was done in the experiment. The value for Φ is selected using an iterative binary search (an interactive, but fast searching algorithm aimed at finding a suitable Φ value) with a threshold of the realized Ω being within 10^-4^ of the desired value. For efficiency, we limit the search over the range Φ = 0-100, as this ranges from neutral selection (Φ=0) to an extremely strong selective pressure (Φ=100) where all cells less (greater) than the mean would die under large (small) selection.

While Malerba et al. (2018) updated centrifugation times and speeds to try and maintain their desired selection percentage (10%), this was not done per-selection event but rather periodically throughout the experiment. As there was not clear documentation on when these adjustments were made, in the model we opted to adjust the value of Φ at each selection event. As adaptation moves cell size in the direction of the selective pressure, the efficacy of the centrifugation protocol is diminished over time thus the realized selective pressure over the course of the experiment was most likely substantially less than the desired 10%. In fact, we found that a substantially lower value of Ω was required to achieve the observed evolutionary trajectories; here Ω= 0.2% for large trajectories and Ω = -3.25% for small lineages.

To avoid any impact that centrifugation might have on the experiment, all lineages including the controls were centrifuged at high speed following selection, which should theoretically pellet all cells. However, this appears to have exerted a slightly positive size selective pressure on the control lineages (all control lineages increase in size across evolutionary trajectories, Figure 3, Supplementary Figure 1). To replicate this process, we also apply a small positive size selection to all model lineages following selection.

To allow for full comparability to the observed evolutionary trajectories from Malerba et al. (2018), we ran the model simulations for two phases mimicking the protocol of Malerba et al. In phase a, we applied size selection, culture dilution, and transfer to new media in the model every 3 to 4 days. After day 900, cultures continued to be diluted and transferred to new media on the same 3 to 4 day interval but selection took place only every 7 days (phase b) (Malerba et al. 2020). Since the frequency of selection was reduced in phase b, in general a stronger selective pressure was needed to maintain size selection, consistent with what was needed experimentally. All values are provided in Supplementary Table 2.

### Phenotype validation simulations

We experimentally assessed the phenotypes of 18 lineages after ~2500 generations of size selection (see Methods below). We compare this experimental phenotype data to the model evolved phenotype in ecoTRACE by running an additional set of model simulations that replicated the experimental set-up. Specifically, we ran a 64 day simulation that consisted of an initial 42 day interim period and then the 22 day phenotyping experiment. The interim period occurred after the evolution experiment before the phenotyping experiment during which the cultures were transferred from the Marshall Lab and acclimated to growth in the Levine Lab. During this period, the cells were transferred every 14 days with a dilution to 20,000 cells mL^-1^ and replenishment of NO3 to 750 μmol L^-1^. While the nitrate concentration in the media was 880 μmol L^-1^, in the simulations we adjusted the modeled dilution concentration to 750 μmol L^-1^ to match the observed nitrate concentrations and drawdown rates in the cultures between days 5-10. Following the interim growth period, the phenotyping experiment was conducted during which the population was diluted to 20,000 cells mL^-1^, transferred to fresh media (NO_3_ = 750 μmol L^-1^) and grown for 22 days without dilution. The entire simulation was done using a 12:12 hr light:dark, cycle with a light intensity of 150 W m^-2^.

### Multi-stressor simulations

Models were initialized with an initial base cell volume of 150 μm^3^, and a 0.1 ExpRate. All model simulations started with a 1,000-day spin-up to allow the population to reach a stable state before the experimental treatments began. In the spinup, nutrients were replenished every 3 days to 880 μmol L^-1^ NO_3_, light was provided on a 12:12 light:day cycle at 150 W m^-2^ irradiance, and diluted back to 30,000 cells during nutrient replenishment. 50 spinup runs were generated, which in turn, were used to initialize each of the 50 replicate runs per treatment. Thus, for each treatment (i.e. high nutrients, low nutrients, pulsed nutrients, etc.), the same 50 initial populations were used as a starting point for each of the stress simulations.

After the 1,000 day spin-up, the populations were allowed to adapt to the treatment condition over 5,000 days of selection. Populations were diluted back to 30,000 cells coinciding with nutrient replenishment (dependent on the treatment, ranging from 1 hour to 100 days). For nutrient fluctuation simulations, an array of simulations were run across two levels of nutrient replenishment (10 and 880 μmol L^-1^ NO_3_), over 5 intervals of replenishment (1 hour, 5-, 10-, 50-, and 100-days). For all the nutrient fluctuation simulations, light was kept at a 12:12 day:night cycle at 150 W m^-2^ irradiance.

For light availability simulations, an array of simulations were run across three light levels (15, 75, and 150 W m^-2^) and four day:night cycles (6:18, 12:12, 18:6, 24:0, Supplementary Figure 5). Nutrients were replenished hourly to 880 μmol L^-1^ NO_3_ and populations were diluted back to 30,000 cells each hour. This time frame maintained the treatment in near-chemostat conditions while allowing super-individuals to create new cells, leading to new mutations between dilutions.

For the combined multi-stressor light-nutrient array, we focused on high nutrient replenishment (880 μmol L^-1^ NO_3_) at 5-, 10-, 50-, and 100-day intervals. This was done over both high (150 W m^-2^) and low (15 W m^-2^) light levels and with all four day:night cycles (6:18, 12:12, 18:6, 24:0) for a total of 1,200 simulations (24 conditions x 50 replicates)

### 5.5 D. tertiolecta Physiology measurements

To validate our model against the evolved phenotypes observed following artificial size-selection experiments, we quantified multiple traits of evolved *D. tertiolecta* lineages across the growth curve. Specifically, we measured cell counts via flow cytometry, cell volume via microscopy, and chlorophyll via fluorometry daily. We also quantified carbon, nitrogen and media nitrate and nitrite at seven time-points over the growth curve (days 3, 5, 6, 7, 8, 9, and 12). Finally, we analyzed cellular carbohydrate content at 8 time-points for large and control lineages, and 4 time-points for the small lineages (Supplementary Figure 8).

#### Culture conditions and maintenance

The 18 evolved lineages were acquired from the Marshall Lab at Monash University on 14 May 2026, 6 per size selection treatment (control, small, and large). Cultures were maintained in f/2 medium - Si at 21 °C and under a 12:12 day:night cycle at a light intensity of 180 umol photon m^- 2^ s^-1^. We followed the weekly size selection protocol currently applied in the Marshall Lab for 2-4 weeks to acclimate the cells. Specifically, selection on the small lineages was conducted by centrifuging the cells at 950 rpm for 5 minutes and retaining the supernatant. Selection for the large lineages was conducted by centrifuging at 380 rpm for 4 minutes and retaining the pellet. The control lineages were not selected for cell size. After size selection, the small and control lineages were centrifuged at 2300 rpm for 8 minutes to pellet the cells. The large lineages were centrifuged at 950 rpm for 5 minutes. The pellets were then resuspended in 20 mL of fresh media. After the acclimation period, all lineages were inoculated into 200 mL of fresh media at an estimated starting density of 5,000 cells mL^-1^.

#### Flow Cytometry Analyses

We measured cell densities for all lineages of *D. tertiolecta* daily via flow cytometry on an Accuri C6 Plus (BD). Specifically, cell numbers were enumerated based on chlorophyll autofluorescence detected in the red fluorescence channel (FL3).

We also assessed the relative neutral lipid content in the cells using the fluorescent stain BODIPY™ 505/515 (Thermo Fisher, MA, USA) following the protocol described in Argyle et al. 2021. Specifically, samples were loaded into a 96 well plate and an aliquot was run to quantify the background fluorescence of the cells. The remaining sample was then stained with BODIPY at a final concentration of 2 μg mL^−1^, incubated for 40 minutes, and re-run on the flow cytometer. Neutral lipid content was defined as the difference in median fluorescence per cell between the pre- and post-stained samples.

#### Cell Volume Microscopy

Cell volumes were quantified every sampling day (except day 10 for control and large lineages and day 11 for small lineages) via microscopy. Samples were preserved in Lugol with a final concentration of 1% and stored in the dark until quantified. Cell dimensions were quantified using a Nikon Eclipse Ti2-E inverted microscope with a 10x Nikon Plan Apo DIC air objective (NA 0.45) and with an additional 1.5x, combined 15x net magnification. An Orca Fusion C14440 digital camera and LED light source (Nikon D-LEDI) was used to take images.

Following image capture, brightfield images were segmented via a global threshold binarization approach. Individual segmented objects were then measured for cross-sectional area and shape. Following the protocol of Malerba et al. (2018), cell volume was estimated based on a prolate spheroid using the equation in (Sun and Liu 2003):

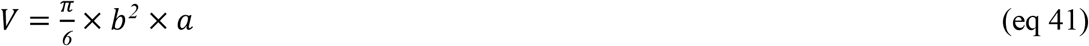

where b = minimum axis length and a = the maximum axis length.

Image processing scripts were used to automate the identification of cells from the background field. Based on differences in morphology for the small, control, and large lineages, lineage specific parameters (circularity, solidity, and minimum and maximum axis length) were used to correctly identify cells (Supplementary Table 4). Cell size was quantified for a minimum of 100 cells per time point per lineage.

#### Chlorophyll Measurements

Chlorophyll concentration was determined fluorometrically using a Trilogy fluorometer with a 460 nm LED module (Turner Designs). A 100 µL sample volume was filtered onto GF/F filters and extracted overnight in 90% acetone at -20 °C in the dark. The fluorometer was calibrated using chlorophyll standards in 90% acetone (Turner Designs).

#### Cellular Carbon and Nitrogen content

Cellular carbon and nitrogen content were measured using an elemental analyzer (Model CEC 440HA; Exeter Analytical) at the UCSB Marine Science Analytical Lab. Samples were collected onto pre-combusted GF/F filters (450°C for 5 hours). We started by filtering 10 mL of culture to ensure sufficient biomass and decreased the working volumes as our cultures grew denser to a minimum of 2 mL by the end of the experiment. The filters were then placed in pre-combusted vials and acidified overnight in the fume hood to remove inorganic substrates following the protocol described in Lepori-Bui et al. 2022. Specifically, un-capped vials were placed in a covered glass tray next to a 100mL beaker containing 75 mL of undiluted HCl for 16 hours. Vials were then placed in a drying oven at 60°C for 2hrs without caps and then for 22hrs with loose caps. Total carbon and nitrogen content was analysed at the UCSB Marine Science Analytical Lab using an Automated Organic Elemental Analyzer (Dumas combustion method, Control Equipment Corp. CEC 440HA).

#### Carbohydrate Analysis

The carbohydrate content of the cells was determined following the protocol of Hu and Finkel (2025). Samples were collected onto combusted GF/F filters under gentle vacuum (130 mmHg). Filters were transferred to cryogenic vials, and immediately flash-frozen in liquid nitrogen before placed at -80 °C until analysis.

#### Media composition

The filtrate from the carbohydrate analysis was collected into combusted 16 mL glass vials for dissolved nutrient analysis. After collection, vials were capped and stored at −20 °C until analysis. Samples were then analyzed for nitrate and nitrite concentration, concentration range 0.1 -300 μM at the UCSB Marine Science Analytical Lab using Flow Injection Analysis (Seal Analytical: SEAL AA500 AutoAnalyzer).

#### Data interpolation

To calculate the daily rate of change in cell volume we used a loess regression fit (R 4.2.2 stats package) against measured values for cell volume. From this smoothed fit we then calculated the rate of change in cell volume per day.

## Supplementary Figures & Tables

**Supplementary Figure 1.**
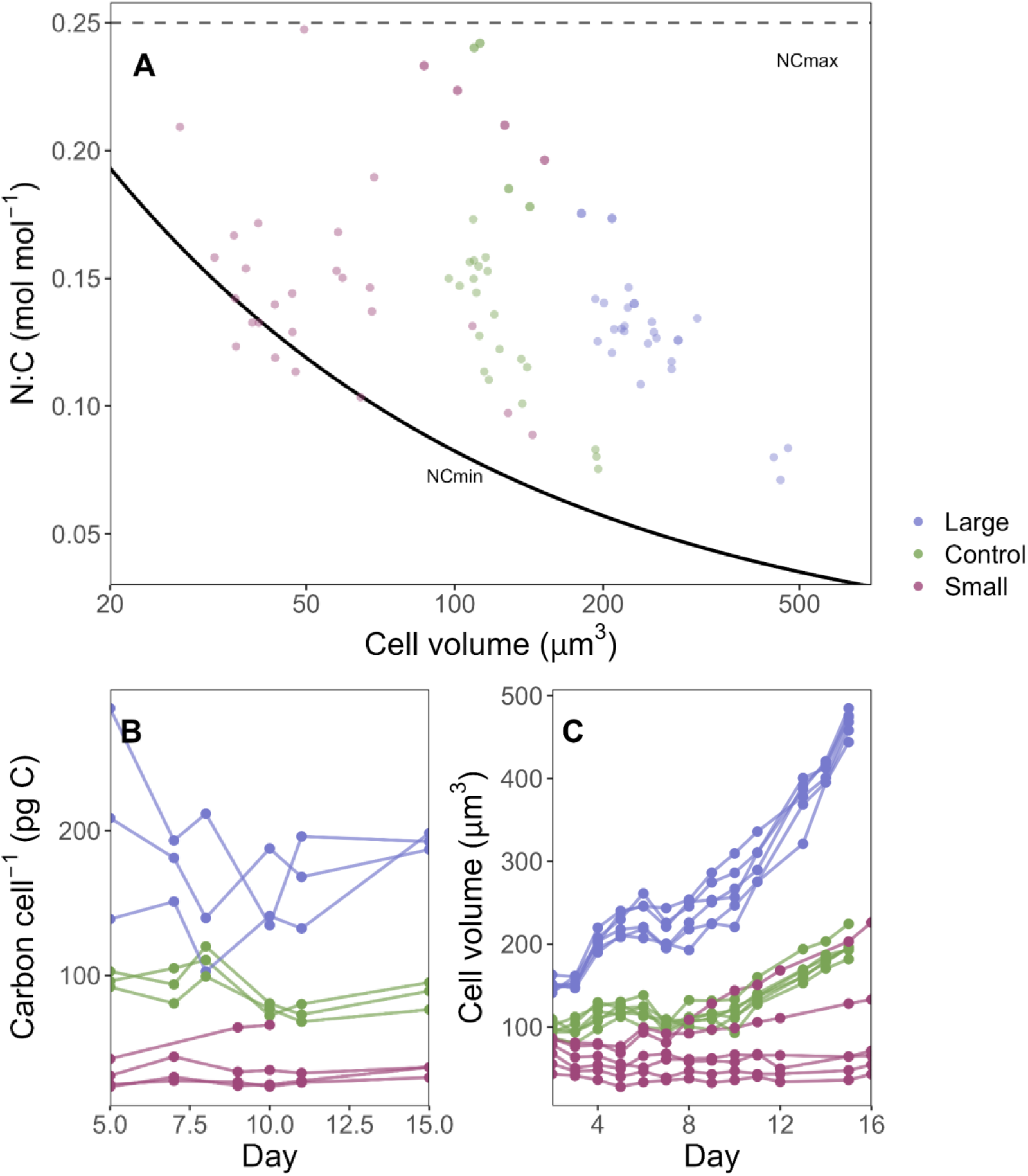
Data for model development and parameter choices. **A**. Cell volume μm^3^ versus Nitrogen:Carbon quota for the three evolutionary lineages (small, control, and large) across the experimental growth curve and during preliminary nutrient limitation experiments. The dashed line shows the NC_max_ value set for the quota model and the solid line shows the NC_min_, first predicted by the proteome model, and later validated with experimental data (eq 18). **B**. Carbon per cell (pg C) over time during experimental growth curves. Connected points represent individual lineages within the experiment. **C**. Cell volume (μm^3^) over time during experimental growth curves. Connected points represent individual lineages within the experiment.

**Supplementary Figure 2:**
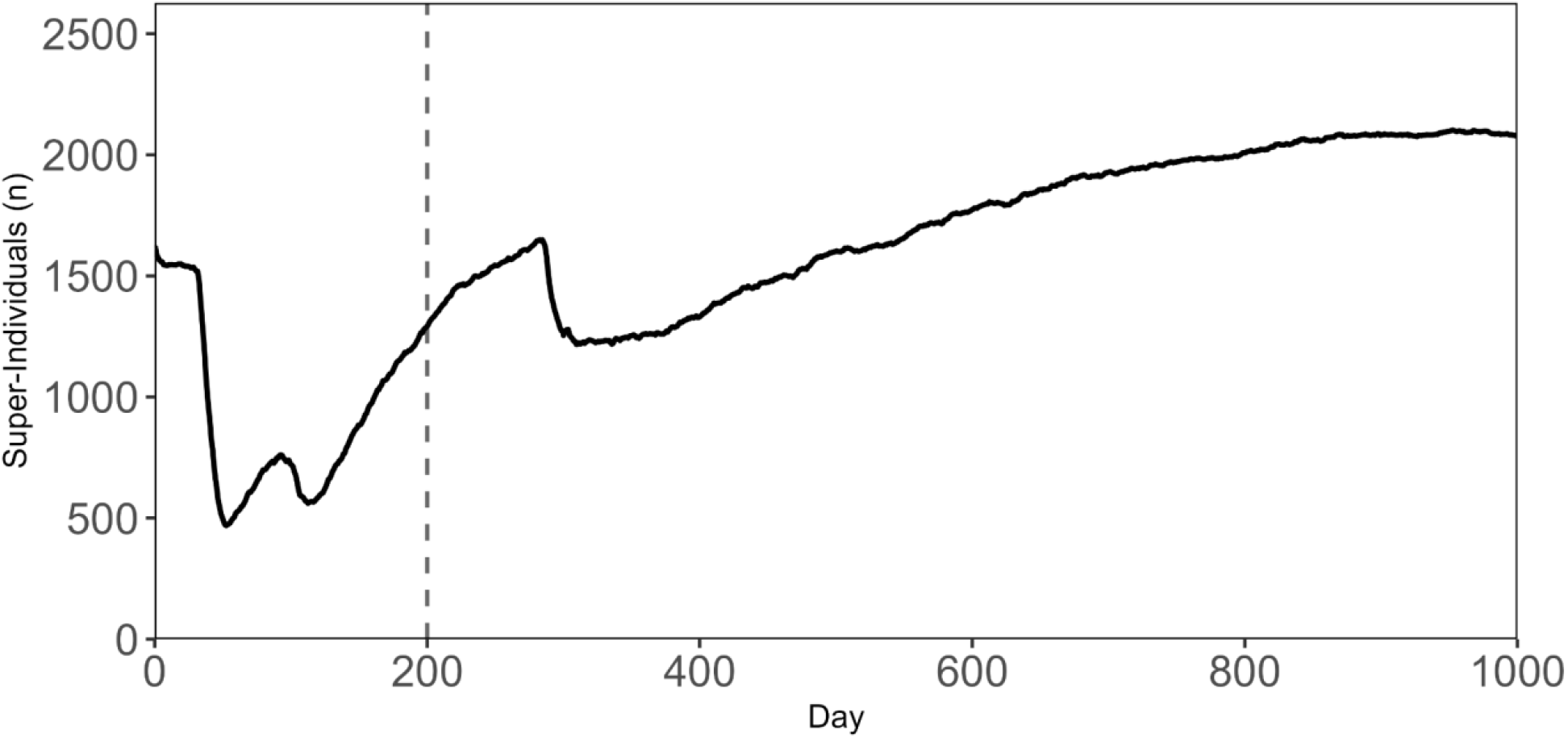
Number of super-individuals over the entire model run presented in (**Figure 1**). The dashed line highlights the first 200 days shown in Figure 2, highlighting initial diversification and a selective sweep.

**Supplementary Figure 3:**
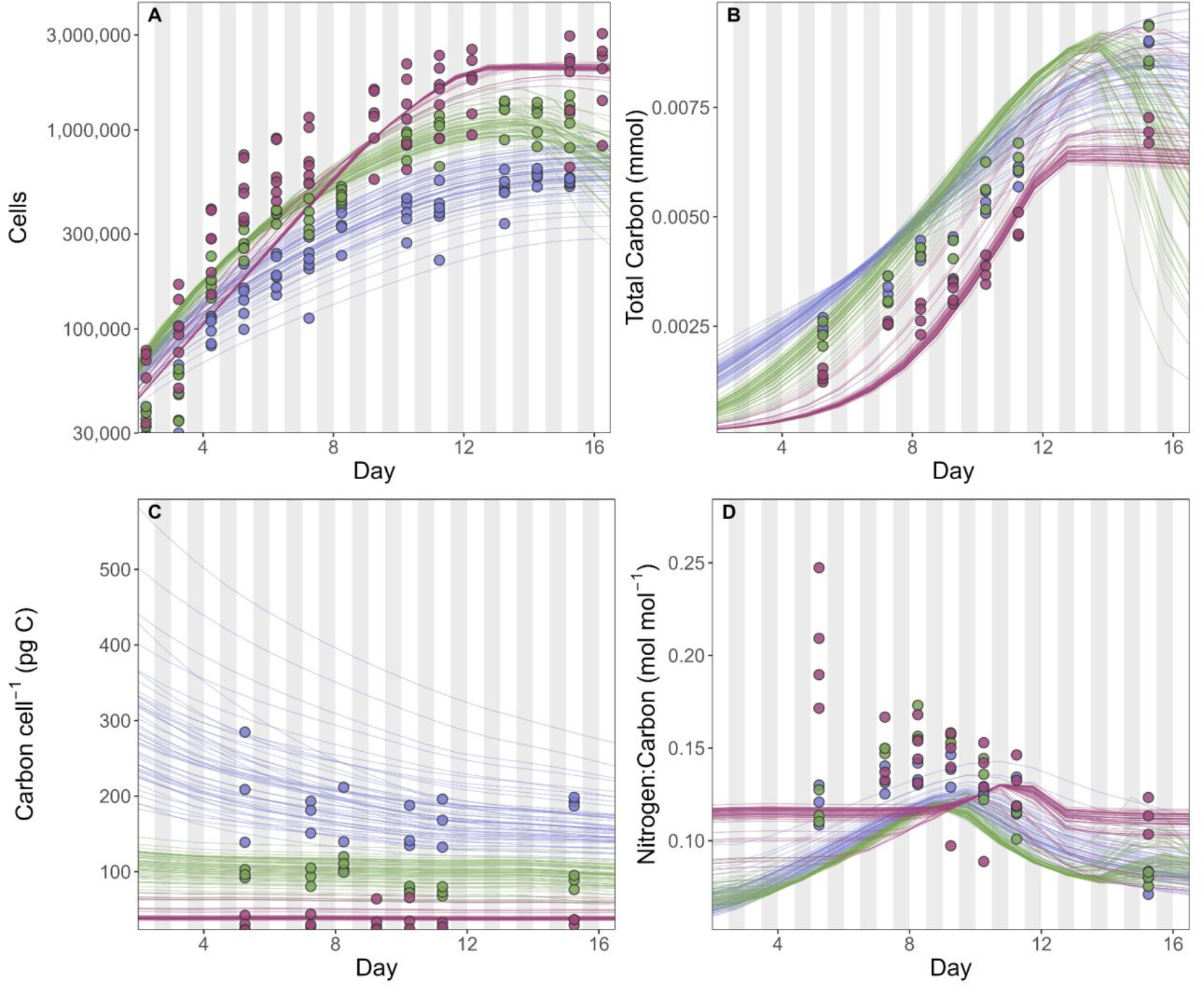
**A**. Comparison between total cells (cells mL^-1^) in the model and growth curve experiments **B**. Comparison between total carbon (mmol C mL^-1^) in the model and growth curve experiments **C**. Comparison between carbon per cell (pg C cell^-1^) in the model and growth curve experiments. **D**. Comparison between Nitrogen:Carbon ratio (mol mol^-1^) in the model and growth curve experiments. Gray shading in **A-D** represents day night cycling.

**Supplementary Figure 4:**
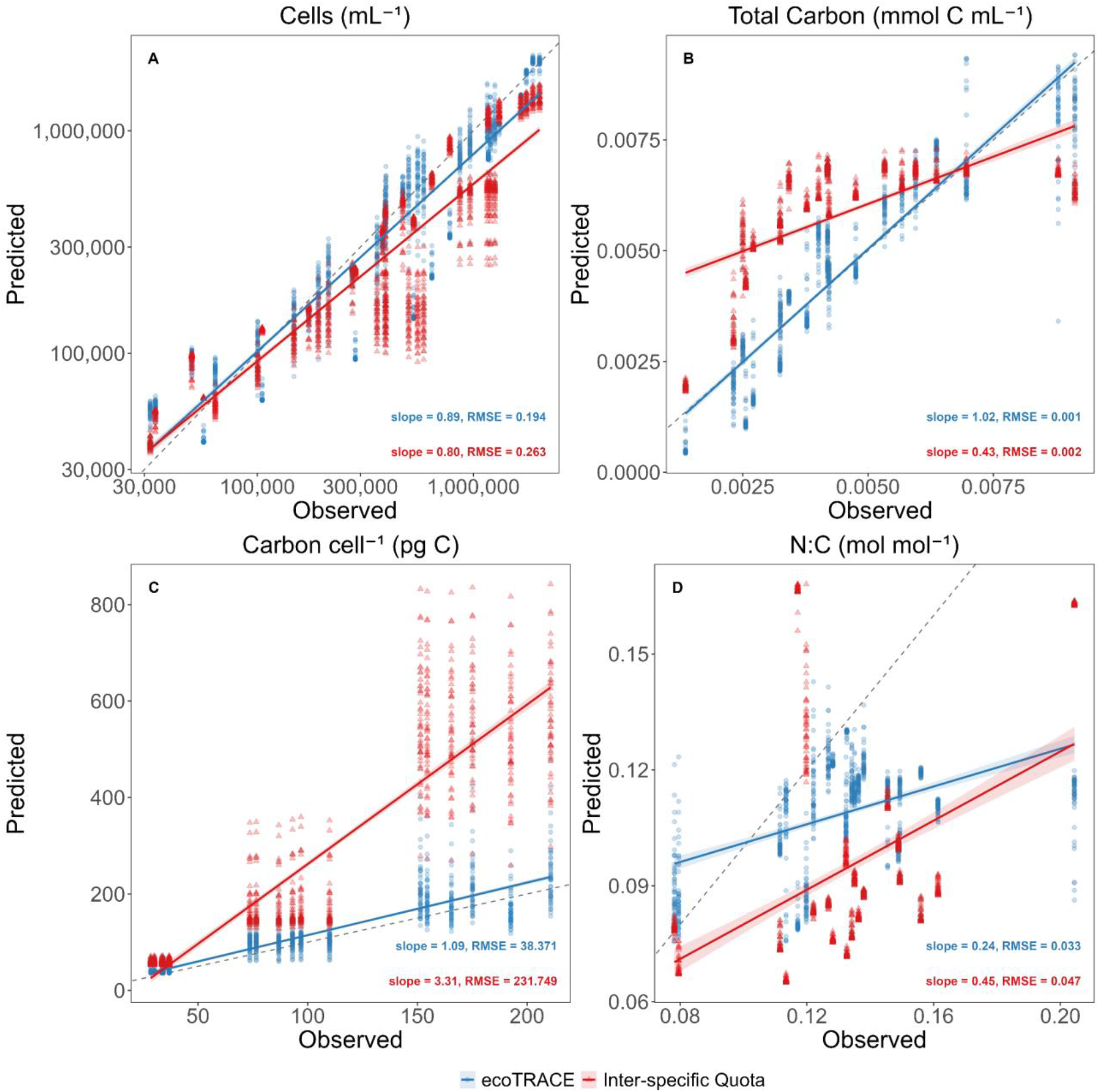
**A-D** Observed (experimental) versus Predicted (either ecoTRACE or inter-specific quota) relationships in key phenotypes. Colored points indicate fit between model and experimental data. Lines represent a generalized linear model across all data per model. Slope and RMSE values are shown per model with colored text to indicate model **A**. Observed vs predicted total Cells (mL^-1^) **B**. Observed vs predicted total carbon (mmol C mL^-1^) **C**. Observed vs predicted carbon per cell (pg C) **D**. Observed vs predicted Nitrogen:Carbon ratio (mol mol^-1^)

**Supplementary Table 1:**
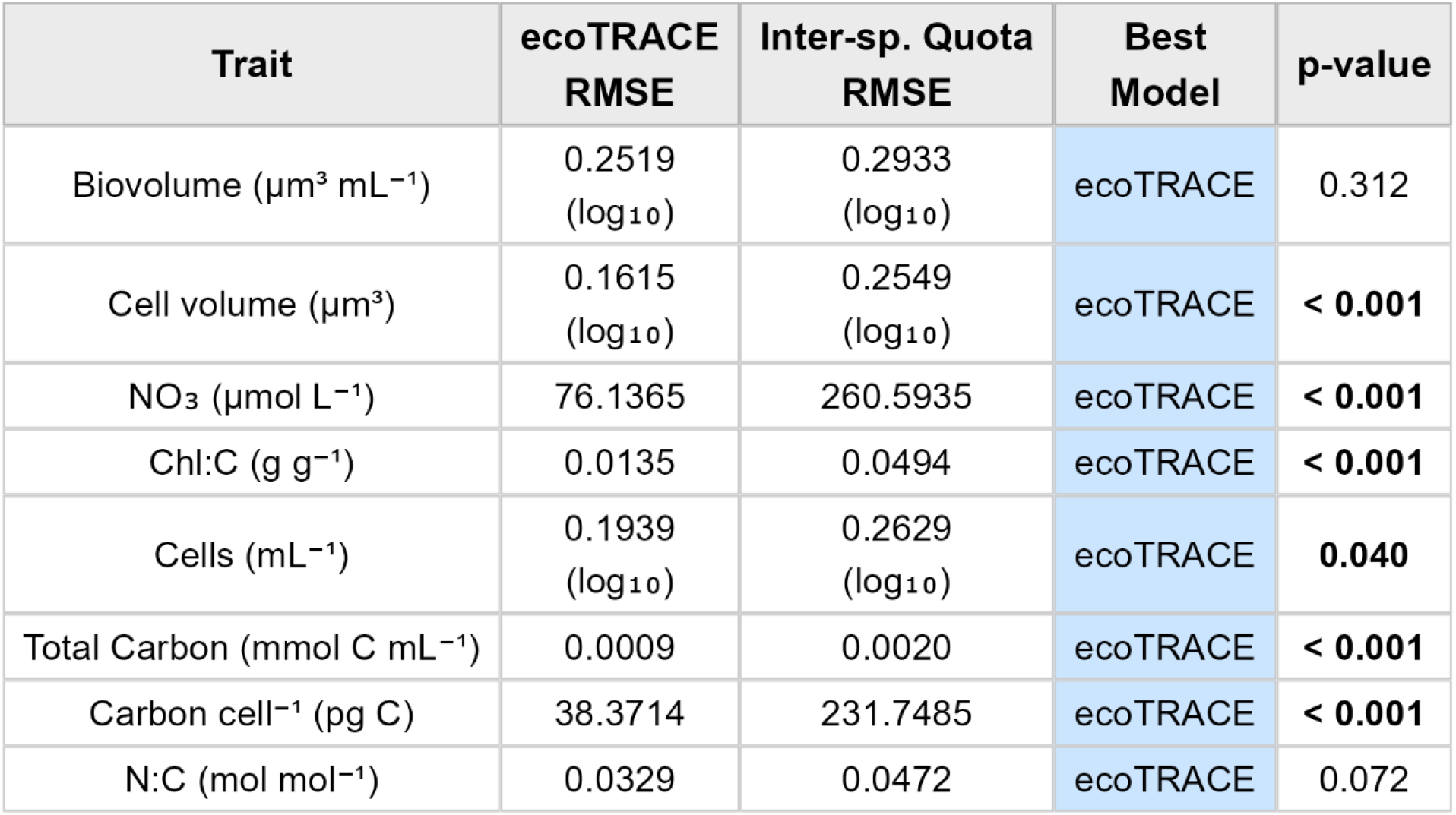
RMSE per trait for both ecoTRACE and inter-specific quota model. P-values are from t-test comparing residuals for ecoTRACE and inter-specific quota model to indicate whether residuals are significantly different between the two model runs.

### Light Only Experiment

Within the light-only experiments, we found that cells primarily adapted via allometric aligned responses, with little to no evidence of plasticity, and that these responses were largely determined by total daily light dose (W m^-2^ day^-1^, **Supplementary Figure 5**). Since nutrient availability in these runs was equivalent to a high N chemostat, populations appeared to adapt by balancing the tradeoff between maximizing photosynthetic capacity, driven by chlorophyll concentration, and minimizing self-shading effects at the population level. Under these conditions, plasticity provided little adaptive advantage within the mechanistic constraints of the model and therefore did not emerge over the evolutionary trajectories.

**Supplementary Figure 5:**
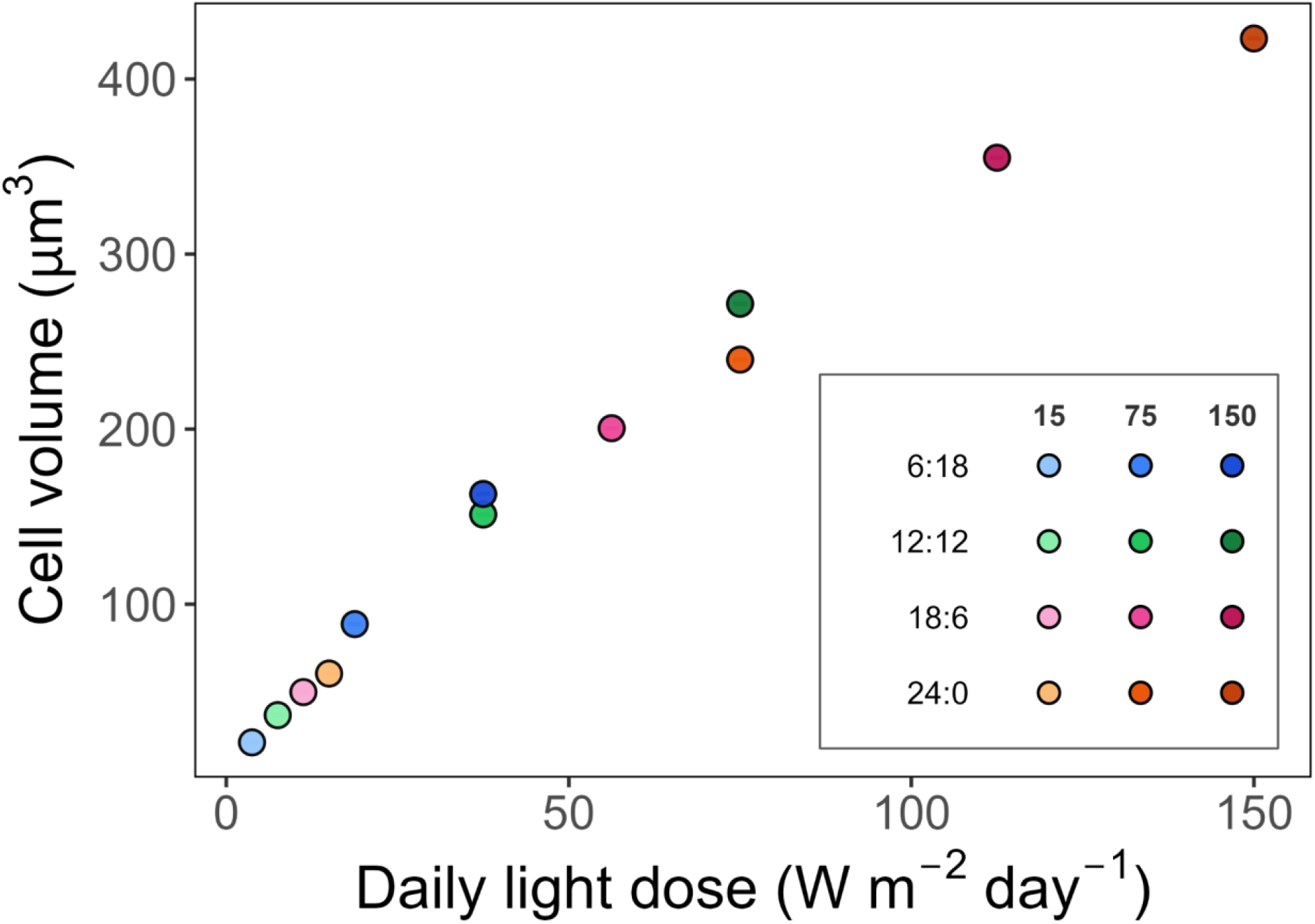
Relationship between daily light dose (W m^-2^ day^-1^) and cell volume (μm^3^) across multiple light-level experiments. Columns in the legend indicate light level (W m^-2^) and rows highlight diel periodicity of day:night cycle (hr:hr).

**Supplementary Figure 6:**
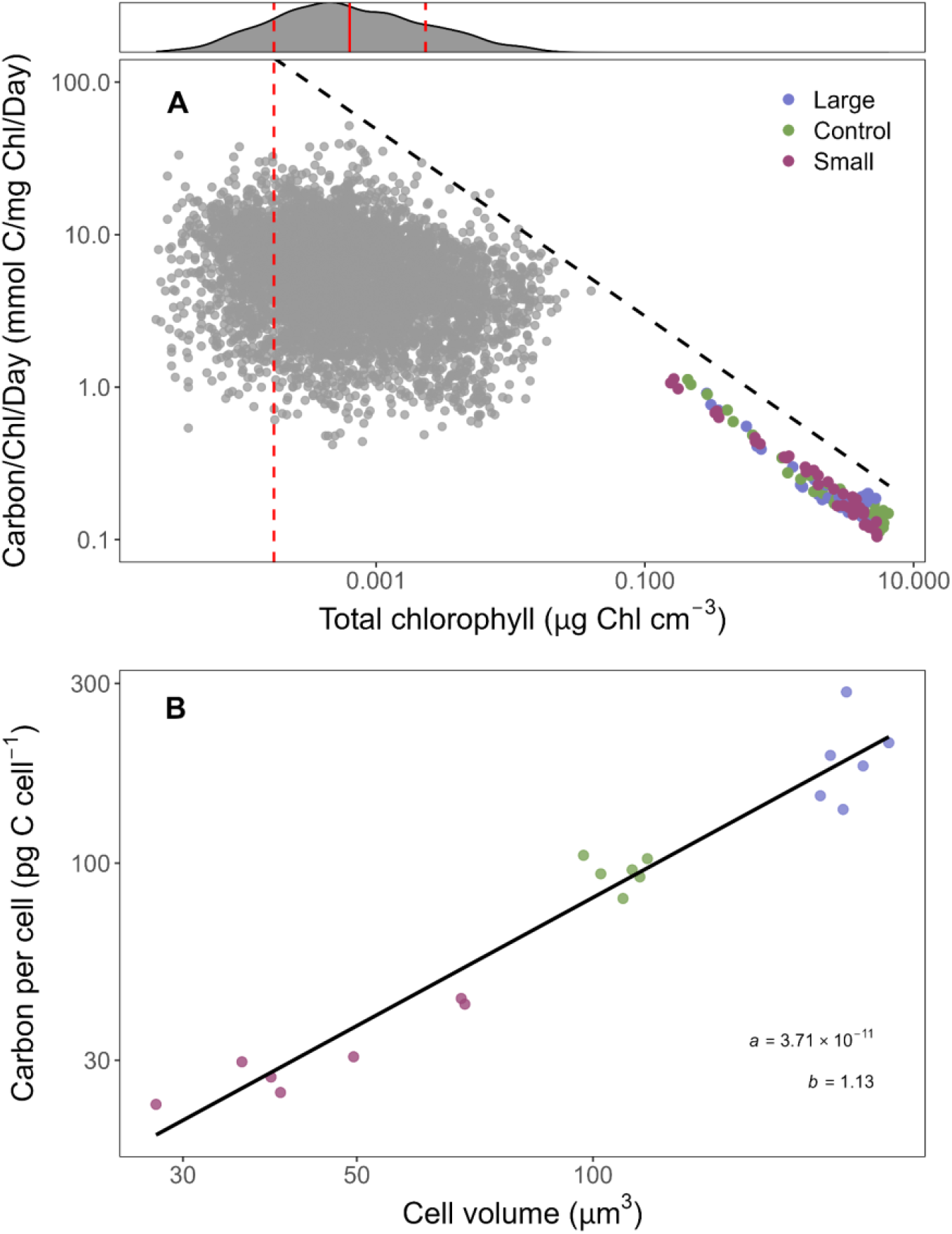
**A**. Relationship between total chlorophyll (μg Chl cm^-3^) and chlorophyll specific carbon fixation rate (Carbon Chl^-1^ Day^-1^, mmol C mg Chl^-1^ day^-1^). Grey points are environmental samples from Bouman et al. (2018). Histogram above panel A shows the distribution of environmental chlorophyll concentration, including mean (red line) and one standard deviation from the mean (dashed red lines). Colored points were estimated from our data on *D. tertiolecta* growth curves. The red dashed line on panel A represents the mean chlorophyll - 1 SD. The dashed black line represents our modeled relationship (eq 21) between chlorophyll concentration and effective chlorophyll specific carbon fixation rate. **B**. Relationship between cell volume and carbon per cell tuned specifically for *D. tertiolecta*. As carbon per cell remains relatively constant per treatment (large, control, small) over time (**Supplementary Figure 1b**), this allometry focused on capturing the carbon per cell during a window where cells are approximately at “base cell volume” (days 3-5 within the growth curve).

**Supplementary Figure 7:**
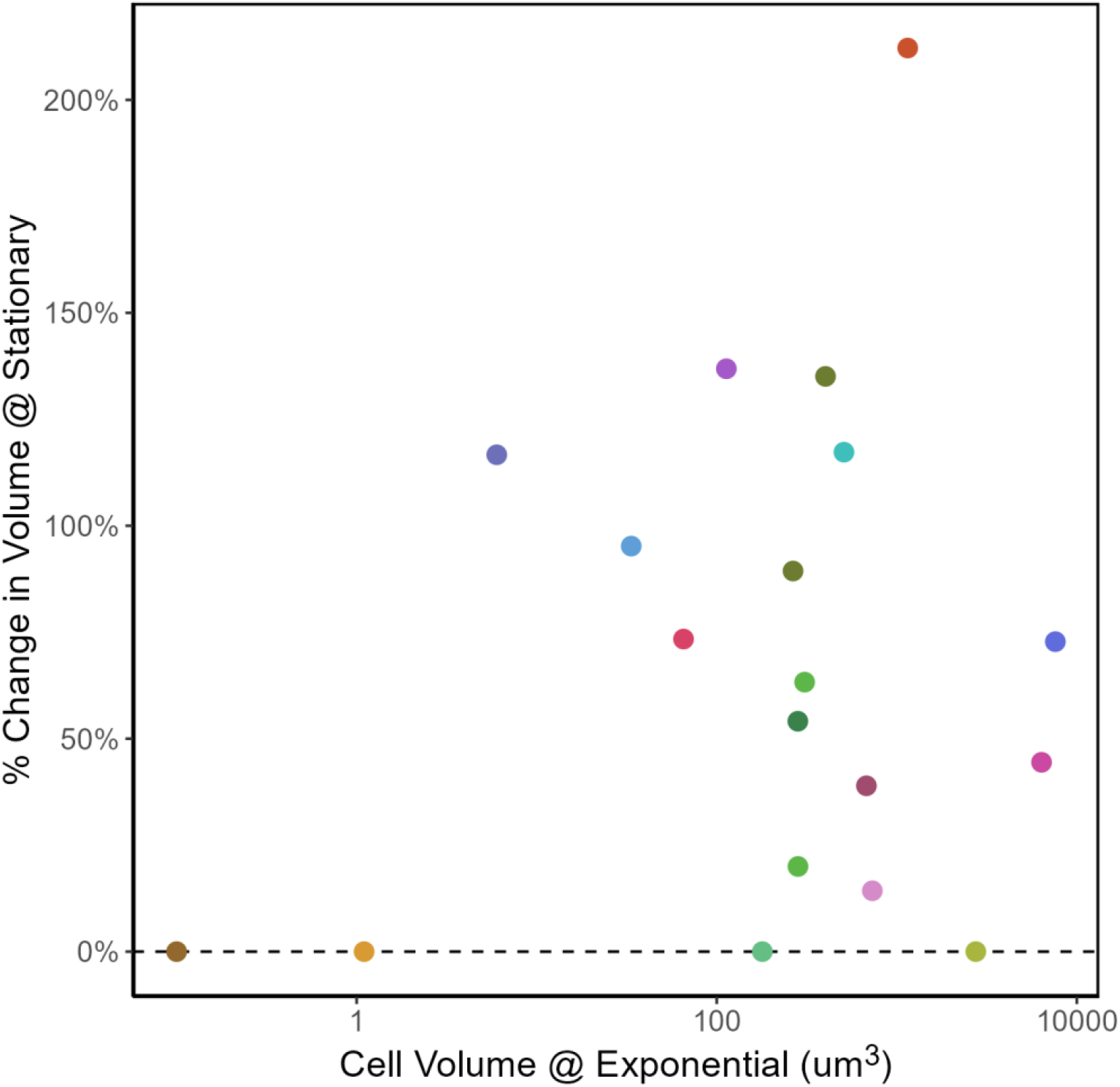
Data from Bigelow collections (Lomas et al. 2024) of culture cell sizes (μm^3^) at exponential versus stationary phase. The x-axis shows the cell size at exponential while the y-axis shows the % change in cell size from exponential to stationary phase. Colors represent the genus of each culture.

**Supplementary Table 2:** Parameters related to selection and centrifugation during the modeled artificial size selection process. Parameters shown for both the ecoTRACE and inter-specific quota models.

| Parameter | Description | ecoTRACE | Standard Quota |
| --- | --- | --- | --- |
| Centrifugation | % increase in volume as a result of "cleaning" centrifugation step | +0.025 | +0.0025 |
| Large sel. phase 1a | Target % volume increase per week, phase 1a | +0.20 | +0.04 |
| Large sel. phase 1b | Target % volume increase per week, phase 1b | +0.225 | +0.04 |
| Small sel. phase 1a | Target % volume decrease per week, phase 1a | -3.25 | -6.50 |
| Small sel. phase 1b | Target % volume decrease per week, phase 1b | -3.25 | -5.50 |
| Adaptive sel. mult. (large) | Multiplier when pop. lags target (large) | 1.30 | 1.30 |
| Adaptive sel. mult. (small) | Multiplier when pop. lags target (small) | 1.075 | 1.075 |

**Supplementary Table 3:** Parameters that were unchanged from Ward and Follows (2016).

| Symbol | Parameter | Value | Units |
| --- | --- | --- | --- |
| **Photosynthesis** |  |  |  |
| $p_a$ | Unimodal $P_{\max,C}$ constant a | 3.08 | — |
| $p_b$ | Unimodal $P_{\max,C}$ constant b | 5.00 | — |
| $p_c$ | Unimodal $P_{\max,C}$ constant c | -3.80 | — |
| $\alpha$ | Initial slope of P-I curve | $3.83 \times 10^{-7}$ | $\text{mmol C (mg Chl)}^{-1} (\mu\text{Ein m}^{-2} \text{s}^{-1})^{-1}$ |
| $h$ | $Q_{\text{stat},N}$ exponent | 0.1 | — |
| **Quotas** |  |  |  |
| $NC_{\min}$ | Minimum N:C quota | 0.05 | $\text{mol N (mol C)}^{-1}$ |
| $NC_{\max}$ | Maximum N:C quota | 0.17 | $\text{mol N (mol C)}^{-1}$ |
| $ChlN^{\max}$ | Maximum Chl:N synthesis rate | 3.0 | $\text{mg Chl (mmol N)}^{-1}$ |
| **N Uptake** |  |  |  |
| $a_{NO_3}$ | $NO_3$ $V_{\max}$ allometry coefficient | 0.44 | $\text{mmol N (mmol C)}^{-1} \text{d}^{-1}$ |
| $a_{NH_4}$ | $NH_4$ $V_{\max}$ allometry coefficient | 0.22 | $\text{mmol N (mmol C)}^{-1} \text{d}^{-1}$ |
| $b_V$ | $V_{\max}$ size exponent | -0.12 | — |
| $k_{NO_3}$ | $NO_3$ half-saturation coefficient | $0.14 \times 10^{-6}$ | $\text{mol L}^{-1}$ |
| $k_{NH_4}$ | $NH_4$ half-saturation coefficient | $0.07 \times 10^{-6}$ | $\text{mol L}^{-1}$ |
| $b_k$ | Half-saturation size exponent | 0.33 | — |
| $\lambda_{NH_4}$ | $NH_4$ inhibition of $NO_3$ uptake | $4.6 \times 10^6$ | $(\text{mol L}^{-1})^{-1}$ |
| **Growth** |  |  |  |
| $\beta_C$ | Biosynthesis cost | 2.33 | $\text{mmol C (mmol N)}^{-1}$ |
| $m$ | Linear mortality rate | 0.05 | $\text{d}^{-1}$ |
| **Temperature** |  |  |  |
| $A$ | Arrhenius activation parameter | -4000 | K |
| $T_0$ | Reference temperature | 25 | $^{\circ}\text{C}$ |

**Supplementary Table 4:** Parameters for the identification of cells via microscopy.

| Treatment | Min. Circularity | Min. Solidity | Minor Axis Length (μm) | Major Axis Length (μm) |
| --- | --- | --- | --- | --- |
| Small | 0.654 | 0.898 | 2.50 | 16.72 |
| Control | 0.799 | 0.949 | 2.50 | 19.80 |
| Large | 0.836 | 0.940 | 2.50 | 25.08 |

**Supplementary Figure 8.**
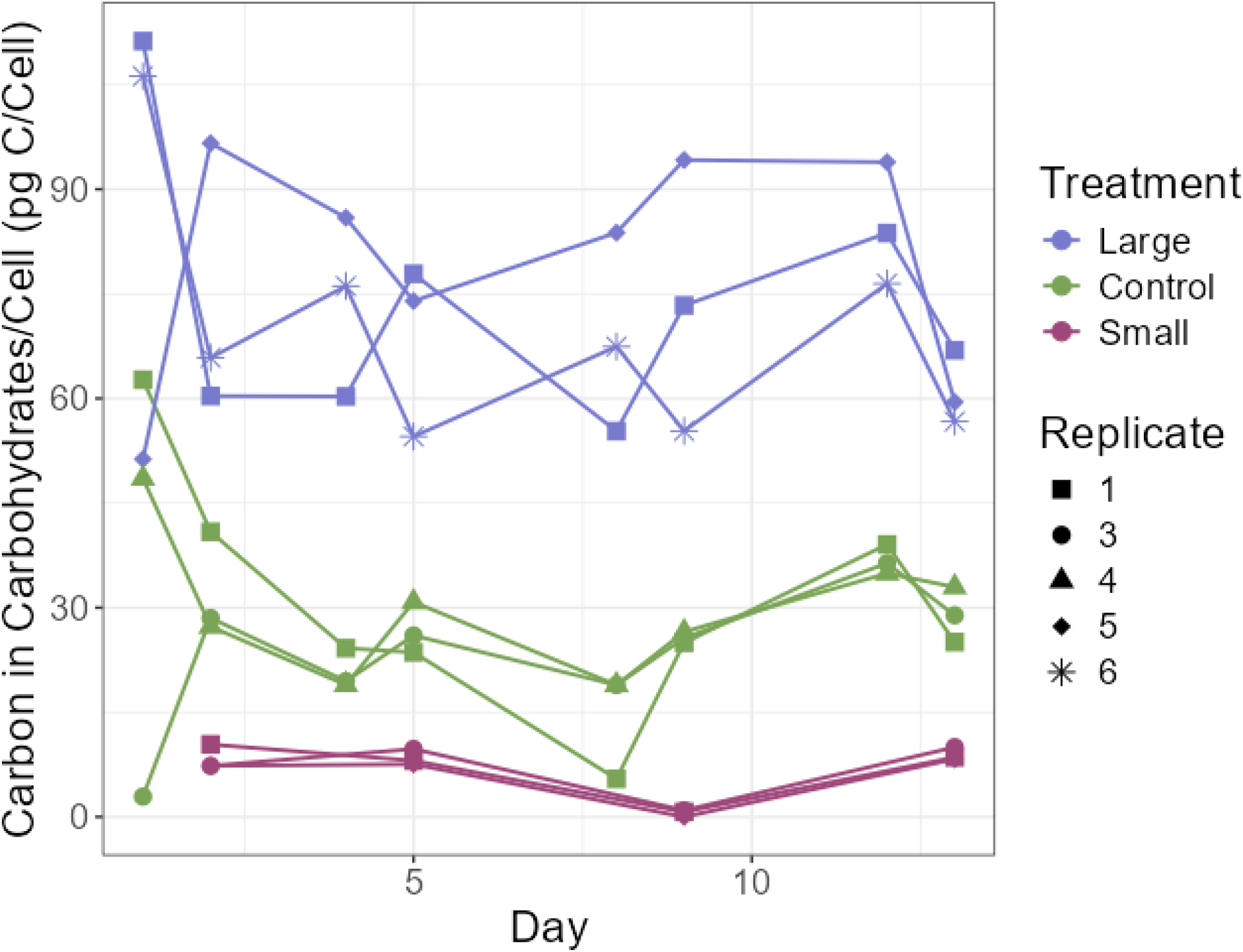
Additional carbohydrate data from growth curve experiment. Carbon in carbohydrates per cell (pg C) over time during experimental growth curves. Connected points represent individual lineages within the experiment.

## References

Anderson, Stephanie I., Clara Fronda, Andrew D. Barton, Sophie Clayton, Tatiana A. Rynearson, and Stephanie Dutkiewicz. “Phytoplankton thermal trait parameterization alters community structure and biogeochemical processes in a modeled ocean.” Global Change Biology 30, no. 1 (2024): 17093.

Agrawal, Anurag A. “A scale-dependent framework for trade-offs, syndromes, and specialization in organismal biology.” Ecology 101, no. 2 (2020): e02924.

Argyle, Phoebe A., Nathan G. Walworth, Jana Hinners, Sinéad Collins, Naomi M. Levine, and Martina A. Doblin. “Multivariate trait analysis reveals diatom plasticity constrained to a reduced set of biological axes.” ISME communications 1, no. 1 (2021): 59.

Aumont, Olivier, Christian Éthé, Alessandro Tagliabue, Laurent Bopp, and Marion Gehlen. “PISCES-v2: an ocean biogeochemical model for carbon and ecosystem studies.” Geoscientific Model Development Discussions 8, no. 2 (2015): 1375–1509.

Benthuysen, Jessica A., Eric CJ Oliver, Ming Feng, and Andrew G. Marshall. “Extreme marine warming across tropical Australia during austral summer 2015–2016.” Journal of Geophysical Research: Oceans 123, no. 2 (2018): 1301–1326.

Bopp, Laurent, Laure Resplandy, James C. Orr, Scott C. Doney, John P. Dunne, M. Gehlen, P. Halloran et al. “Multiple stressors of ocean ecosystems in the 21st century: projections with CMIP5 models.” Biogeosciences 10, no. 10 (2013): 6225–6245.

Bouman, Heather A., Trevor Platt, Martina Doblin, Francisco G. Figueiras, Kristinn Gudmundsson, Hafsteinn G. Gudfinnsson, Bangqin Huang et al. “Photosynthesis–irradiance parameters of marine phytoplankton: synthesis of a global data set.” Earth System Science Data 10, no. 1 (2018): 251–266.

Brandenburg, Karen M., Sylke Wohlrab, Uwe John, Anke Kremp, Jacqueline Jerney, Bernd Krock, and Dedmer B. Van de Waal. “Intraspecific trait variation and trade-offs within and across populations of a toxic dinoflagellate.” Ecology Letters 21, no. 10 (2018): 1561–1571.

Cai, Wenju, Benjamin Ng, Guojian Wang, Agus Santoso, Lixin Wu, and Kai Yang. “Increased ENSO sea surface temperature variability under four IPCC emission scenarios.” Nature Climate Change 12, no. 3 (2022): 228–231.

Cheng, Chuankai, Brittany D. Bennett, Pratixa Savalia, Hasti Asrari, Carmen Biel, Kate A. Evans, Rui Tang, and J. Cameron Thrash. “Cell cycle dysregulation of globally important SAR11 bacteria resulting from environmental perturbation.” Nature Microbiology (2026): 1–15.

Collins, Sinéad, Harriet Whittaker, and Mridul K. Thomas. “The need for unrealistic experiments in global change biology.” Current Opinion in Microbiology 68 (2022): 102151.

Droop, Michaël R. “Vitamin B12 and marine ecology. IV. The kinetics of uptake, growth and inhibition in Monochrysis lutheri.” Journal of the Marine Biological Association of the United Kingdom 48, no. 3 (1968): 689–733.

Edwards, Kyle F., Christopher A. Klausmeier, and Elena Litchman. “Nutrient utilization traits of phytoplankton: ecological archives E096-202.” Ecology 96, no. 8 (2015): 2311–2311.

Edwards, Kyle F., Mridul K. Thomas, Christopher A. Klausmeier, and Elena Litchman. “Light and growth in marine phytoplankton: allometric, taxonomic, and environmental variation.” Limnology and Oceanography 60, no. 2 (2015): 540–552.

Follows, Michael J., Stephanie Dutkiewicz, Scott Grant, and Sallie W. Chisholm. “Emergent biogeography of microbial communities in a model ocean.” science 315, no. 5820 (2007): 1843–1846.

Geider, R. J., H. L. MacIntyre, and T. M. Kana. “Dynamic model of phytoplankton growth and acclimation: responses of the balanced growth rate and the chlorophyll a: carbon ratio to light, nutrient-limitation and temperature.” Marine Ecology Progress Series 148 (1997): 187–200.

Henson, Stephanie A., B. B. Cael, Stephanie R. Allen, and Stephanie Dutkiewicz. “Future phytoplankton diversity in a changing climate.” Nature communications 12, no. 1 (2021): 5372.

Hillebrand, Helmut, Esteban Acevedo-Trejos, Stefanie D. Moorthi, Alexey Ryabov, Maren Striebel, Patrick K. Thomas, and Marie-Luise Schneider. “Cell size as driver and sentinel of phytoplankton community structure and functioning.” Functional Ecology 36, no. 2 (2022): 276–293.

Hu, Ying-Yu, and Zoe V. Finkel. “Total particulate carbohydrate from microalgae.” (2025).

Leles, Suzana G., and Naomi M. Levine. “Mechanistic constraints on the trade-off between photosynthesis and respiration in response to warming.” Science Advances 9, no. 35 (2023): eadh8043.

Leles, Suzana G., Lara Breithaupt, Arianna Krinos, Harriet Alexander, Holly V. Moeller, Lana Flanjak, Charlotte Laufkotter, Elena Litchman, María Aranguren-Gassis, and Naomi M. Levine. “New niches for larger phytoplankton in a warmer, more resource-limited ocean.” bioRxiv (2025): 2025–06.

Lepori-Bui, Michelle, Christopher Paight, Ean Eberhard, Conner M. Mertz, and Holly V. Moeller. “Evidence for evolutionary adaptation of mixotrophic nanoflagellates to warmer temperatures.” Global Change Biology 28, no. 23 (2022): 7094–7107.

Levine, Naomi M., Martina A. Doblin, and Sinéad Collins. “Reframing trait trade-offs in marine microbes.” Communications Earth & Environment 5, no. 1 (2024): 219.

Li, Guancheng, Lijing Cheng, Jiang Zhu, Kevin E. Trenberth, Michael E. Mann, and John P. Abraham. “Increasing ocean stratification over the past half-century.” Nature Climate Change 10, no. 12 (2020): 1116–1123.

Li, William KW. “Temperature adaptation in phytoplankton: cellular and photosynthetic characteristics.” Primary productivity in the sea (1980): 259–279.

Lomas, M. W., A. R. Neeley, R. Vandermeulen, A. Mannino, C. Thomas, M. G. Novak, and S. A. Freeman. Phytoplankton optical fingerprint libraries for development of phytoplankton ocean color satellite products, Sci. Data, 11, 168. 2024.

Malerba, Martino E., Craig R. White, and Dustin J. Marshall. “Eco-energetic consequences of evolutionary shifts in body size.” Ecology Letters 21, no. 1 (2018): 54–62.

Malerba, Martino E., Maria M. Palacios, Yussi M. Palacios Delgado, John Beardall, and Dustin J. Marshall. “Cell size, photosynthesis and the package effect: an artificial selection approach.” New Phytologist 219, no. 1 (2018): 449–461.

Malerba, Martino E., and Dustin J. Marshall. “Size-abundance rules? Evolution changes scaling relationships between size, metabolism and demography.” Ecology Letters 22, no. 9 (2019): 1407–1416.

Malerba, Martino E., Giulia Ghedini, and Dustin J. Marshall. “Genome size affects fitness in the eukaryotic alga Dunaliella tertiolecta.” Current Biology 30, no. 17 (2020): 3450–3456.

Malerba, Martino E., and Dustin J. Marshall. “Larger cells have relatively smaller nuclei across the Tree of Life.” Evolution Letters 5, no. 4 (2021): 306–314.

Müller, Oliver, Tommaso Redaelli, Dieter A. Baumgartner, Clara Martínez-Pérez, Francesco Carrara, Johannes M. Keegstra, and Roman Stocker. “Heritable diel energy reserves enhance diatom growth.” bioRxiv (2026): 2026–01.

O’Donnell, Daniel R., Sophia M. Beery, and Elena Litchman. “Temperature-dependent evolution of cell morphology and carbon and nutrient content in a marine diatom.” Limnology and Oceanography 66, no. 12 (2021): 4334–4346.

Peter, Kalista Higini, and Ulrich Sommer. “Phytoplankton cell size reduction in response to warming mediated by nutrient limitation.” PloS one 8, no. 9 (2013): e71528.

Richter, Daniel J., Romain Watteaux, Thomas Vannier, Jade Leconte, Paul Frémont, Gabriel Reygondeau, Nicolas Maillet et al. “Genomic evidence for global ocean plankton biogeography shaped by large-scale current systems.” elife 11 (2022): e78129.

Stock, Charles A., John P. Dunne, and Jasmin G. John. “Global-scale carbon and energy flows through the marine planktonic food web: An analysis with a coupled physical–biological model.” Progress in Oceanography 120 (2014): 1–28.

Sun, Jun, and Dongyan Liu. “Geometric models for calculating cell biovolume and surface area for phytoplankton.” Journal of plankton research 25, no. 11 (2003): 1331–1346.

Walworth, Nathan G., Jana Hinners, Phoebe A. Argyle, Suzana G. Leles, Martina A. Doblin, Sinéad Collins, and Naomi M. Levine. “The evolution of trait correlations constrains phenotypic adaptation to high CO2 in a eukaryotic alga.” Proceedings of the Royal Society B: Biological Sciences 288, no. 1953 (2021): 20210940.

Ward, Ben A., and Michael J. Follows. “Marine mixotrophy increases trophic transfer efficiency, mean organism size, and vertical carbon flux.” Proceedings of the National Academy of Sciences 113, no. 11 (2016): 2958–2963.

Ward, Ben A., Stephanie Dutkiewicz, Oliver Jahn, and Michael J. Follows. “A size-structured food-web model for the global ocean.” Limnology and Oceanography 57, no. 6 (2012): 1877–1891.

